# Sex shapes capillary flows and oxygen extraction in the middle-aged mouse brain

**DOI:** 10.64898/2026.10.12.758495

**Authors:** Eugenio Gutiérrez-Jiménez, Boris Wied, Thomas B. Lindhardt, Signe H. Mikkelsen, Rasmus West Knopper, Christian Stald Skoven, Oliver B. Christensen, Vitalii Dashkovskyi, Filip Mrotek, Peter M. Rasmussen, Maria H. Andresen, Aleksandar Krmpot, Mirna El Khatib, Sergei Vinogradov, Sava Sakadžić, Malene Fjorside Jensen, Isabela Pedroza-Pacheco, Dmitry Postnov, Lin Lin, Jesper Just, Brian Hansen, Leif Østergaard

## Abstract

Sex differences in brain aging are well documented, but their origin within the cerebrovascular hierarchy remains unclear. Here we show that cortical blood flow is higher in female than male mice at 15 months of age, at a point when an earlier sex difference in pulsatility and resistance indices has already converged, pial vessel diameter shows no difference between sexes, and estrogen levels are similar between sexes. Using an indicator-dilution and intravascular oxygen measurements using two-photon microscopy, we found that capillary transit-time heterogeneity (CTH) scales more steeply with mean transit time in females, consistent with a passively regulated capillary network, and that this is accompanied by reduced oxygen extraction fraction (OEF) without deficit in arterial oxygen delivery. Spatial transcriptomics reveals that sex differences are organized into distinct regional programs, most prominently in white matter, where a male-associated myelin and axonogenesis signature accompanies reduced white matter volume in females. These molecular and structural differences coincide with elevated hippocampal microglial reactivity and an altered spatial search strategy in females, without any detectable differences in circulating markers of stress, blood-brain barrier integrity, or immunity. Together, these findings indicate that sex differences in the aging brain are expressed at the level of capillary flow regulation and regional tissue programs rather than through bulk cerebral blood flow alone, and that elevated blood flow in females can mask, rather than reflect, adequate tissue oxygenation.

## Background

Aging is a continuous process that affects all aspects of the body, including the brain, and is a primary risk factor for neurodegenerative disease, such as Alzheimer’s disease (AD)^1^. Although age cannot be modified, 14 other risk factors for dementia have been identified as potentially modifiable, and risk reduction should begin early and continue throughout life^2^. This makes midlife an important window for studying how brain aging begins. Whether sex shapes this process remains debated. Dementia incidence is higher in women than men in some but not all countries, and in one large meta-analysis this difference was explained by longer life expectancy and educational differences, although reduced estrogen after menopause has been proposed as an additional contributor^2^. Studies of cerebral perfusion across the menopause transition have reported lower cerebral blood flow (CBF)^3^, preserved cerebrovascular reactivity^4^, or elevated regional perfusion^5^. Therefore, it remains unclear at which level of the vascular hierarchy sex differences arise in middle age.

CBF, pulsatility index (PI), and resistance index (RI) are only rough predictors of effective brain perfusion. PI reflects vascular tone and compliance, and cross-sectional human data suggest it rises faster with age in women than in men during midlife, largely through carotid stiffening and pulse pressure^6^. In mice, microvascular pulsatility can differ between sexes without a difference in mean perfusion^7^. Oxygen delivery also depends on how blood is distributed across the capillary level, where heterogeneous capillary transit times limit the oxygen that can be extracted at a given flow^8^. Capillary flow is heterogeneous at rest and homogenizes during activation, with mean transit time (MTT) and capillary transit-time heterogeneity (CTH) changing in parallel in passive networks^9^. In contrast, at low flow or during high metabolic demands, active homogenization of capillary flow deviates from this relationship^9,10^. Elevated CTH has been reported in established and prodromal AD^11,12^, and capillary flow disturbances precede memory impairment in a mouse model of cerebral amyloidosis^13^. To our knowledge, whether sex influences capillary flow regulation in middle age has not been examined.

Sex differences in brain aging may also be expressed at the cellular and regional organization of brain tissue. In the mouse hippocampus, microglial gene expression diverges between sexes with age, with females showing enrichment in inflammatory and senescence signatures^14,15^, and spatial transcriptomics of the aging brain has identified white matter as a region of prominent age-related change with a stronger neuroinflammatory response in females^16^. In humans, white matter organization and cortical volume across the menopause transition follow different patterns at every stage of the transition, rather than simple decline^17,18^. Circulating stress hormones are also of interest, since sex-specific associations between cortisol and brain biomarkers have been reported in midlife despite similar cortisol levels in men and women^19^. These observations suggest that sex differences may be concentrated in particular brain regions, where whole-brain measures can miss them, and that mapping gene expression across tissues makes it possible to relate it to vascular function.

In this study, we asked how sex shapes brain aging in middle age, and at what vascular level differences between females and males first appear. We studied male and female mice between 13 and 15 months of age. We combined behavior and blood flow in the living brain with imaging of capillary flow dynamics and oxygen extraction, regional gene expression, and tissue histology. At this age, female mice had higher CBF and more spatial errors. PI, RI differences were transient, and pial vessel diameter did not differ. However, we observed signs of disturbed capillary flow regulation and reduced oxygen extraction, accompanied by differences in regional gene expression. These findings suggest that sex differences in the aging brain are most clearly expressed at the capillary level. At the same time, CBF, PI and RI alone provide an incomplete picture of oxygen supply.

## Methods

### Animal Handling and preparation

The Danish Animal Experiment Inspectorate approved all procedures (Permit 2022-15-0201-01118), in line with the European Council actions for the protection of animals used for scientific purposes and the ARRIVE Criteria.

A total of 12 male and female C57BL/6NRj mice aged 13-15 months were bred at the animal facility at the Department of Biomedicine, Aarhus University, Denmark. At 3 months of age, mice were transferred to the temporary animal facility at the Center for Functionally Integrative Neuroscience (CFIN), where they were housed until the end of the experiments. A set of five 7-months-old female mice with C57BL/6NRj background were selected for plasma analysis of estradiol, but no more analysis were performed. Mice were housed in groups of a maximum of 5 mice per cage in a temperature– and humidity-controlled environment (22-24 °C, 55 ± 10% humidity), a 12-h light/12-h dark cycle (06:00–18:00), with access to food and water *ad libitum*.

At 12 months of age, mice were implanted with a chronic cranial window, head-bar, and neck-collar for awake-restrained imaging using laser speckle contrast imaging (LSCI), two-photon microscopy (2P), and magnetic resonance imaging (MRI) systems, as previously described in a protocol published by our group^20^. Two weeks before surgery, mice were handled daily to reduce stress from procedures and scans and improve postsurgical recovery. To reduce brain edema, mice were administered dexamethasone 4.8 mg/kg (Kalcex, Riga, Latvia) 1 h before anesthesia. Surgical anesthesia was induced with ketamine (87.5 mg/kg) and xylazine (6.25 mg/kg) administered intraperitoneally (i.p.), and topped with 1/3 of the dosage every 30-45 min after checking for reflexes (i.e., tail pinching and pedal reflex). Eye ointment (Viscotears, carbomer 2 mg/g) was applied to each eye to protect against dryness. Subsequently, i.p. injections of ampicillin (200 mg/kg; Stada, Bad Vilbel, Germany) and carprofen (5 mg/kg; Huvepharma, Sofia, Bulgaria) were administered as antibiotic and anti-inflammatory/pain-relief medication, respectively. Mice were securely mounted on a stereotaxic frame (RWD Life Science Co., LTD, Shenzhen, Guangdong, China), and local anesthesia with lidocaine was administered subcutaneously.

Ventilation support during surgical preparation was provided using a mixture of oxygen and medical air (inspired fraction of oxygen [FiO2], 50%) delivered via a face mask. Body temperature was maintained at 37 °C using a heating pad with a feedback-control probe placed in the mouse’s rectum. The mouse’s cranium was exposed after making a small incision in the scalp. The skin was fixed to the skull with cyanoacrylate glue (Loctite Super Glue gel; Loctite Brand, Westlake, Ohio, USA), and grooves were made on the skull (posterior frontal, parietal, and occipital bones) to improve fixation of the dental cement used to maintain the cranial window and the neck holder in place. A cranial window of ≈3 mm in diameter was made centered over the left primary somatosensory barrel cortex brain region representing the whisker C3 (1.5–2 mm posterior, 3 mm mediolateral relative to bregma) using a dental drill equipped with a diamond-cut 0.5-mm burr, and the dura mater was left intact. Once the piece of skull was removed and it was confirmed that no active bleeding was present, the window was covered with an in-house glued glass plug (3 mm in diameter). The plug was secured by applying a thin layer of cyanoacrylate glue gel around the edges. 3D-printed neck-collar and head-bar were centered over the occipital and frontal bones, respectively, and secured with a drop of cyanoacrylate glue. Dental cement (Meliodent; Kulzer GmbH, Hanau, Germany) was applied to the skull along with the 3D-printed pieces to keep them fixed for the remainder of the experiment. Once the dental cement had hardened, a dose of atipamezole hydrochloride 1 mg/kg was administered i.p. to reverse the anesthesia. Mice were maintained on a heating pad with 100% oxygen until they showed signs of voluntary movement. Subsequently, mice were placed in a thermoregulated recovery cage and remained there until complete recovery from anesthesia. Mice were followed during the postsurgical period (≈5 days), providing pain relief with carprofen (10 mg/kg, i.p.) and buprenorphine (Temgesic, 0.06 mg/kg, i.p.; Indivior Europe Limited, Dublin, Ireland), and antibiotic coverage with ampicillin (200 mg/kg; i.p.).

Following the post-surgical period, mice were trained in accordance with in-house protocols for awake imaging with MRI and optical imaging, as previously described^20^. For this study, MRI training was performed before optical imaging training because MRI was considered the most stressful procedure. MRI habituation occurred over 10 consecutive days in a mock MRI environment, as previously described^21^. Training started with 20 min of restraint daily, increasing by 20-min increments until 2.5 h. An MRI scan was performed on day 11, and the mice were allowed to rest for 1 day. After a rest day, mice were habituated for 2 days on the holder for the optical imaging systems and then scanned using LSCI and 2P on subsequent days, with one rest day between scanning sessions.

#### In vivo MRI scans

Scans were performed on a Bruker BioSpec 9.4T (Avance III) small animal MRI system equipped with a 76 mm quadrature transmission volume coil and a four-element rodent brain surface array cryoprobe for reception. The system is equipped with BGA-12HP gradients capable of 650 mT/m (Bruker Biospin, Ettlingen, Germany). Awake mouse imaging was performed with thoroughly habituated mice, as previously described^20^. Briefly, this involved habituation to restraint using a surgically implanted neck-collar attached to a 3D-printed animal bed. Wax earplugs were placed in the mouse’s ears to dampen the sound from the gradient coil. Kwik-cast (WPI, Sarasota, FL, USA) was applied to reduce air between the skull and the coil, then removed after the scanning session. CBF was estimated using pseudo-continuous arterial spin labeling (pCASL) as described in^22^ existing procedures for awake mouse MRI^20,21^. Total scan time for CBF estimation was roughly 1 h, including animal positioning and high-resolution field shimming.

CBF was estimated using in-house MATLAB scripts (MathWorks, Natick, MA, USA)^20,22^. Cortical regions of interest (ROIs) were delineated contralaterally to the cranial window. Pixelwise CBF estimates were then extracted from these ROIs and pooled by group (sex, age) following outlier removal.

#### Laser Speckle Contrast Imaging (LSCI)

A custom LSCI system was used to acquire images to estimate microvascular blood flow index (BFI), pulsatility index (PI), resistance index (RI), and heart rate (HR) as previously described^7^. Incident illumination of the cranial window was provided by a highly coherent, volume holographic grating-stabilized laser diode (785 nm, Thorlabs FPV785P, Newton, NJ, USA), driven by a current controller (LDC205C, Thorlabs) and temperature controller (TED200C, Thorlabs).

Backscattered light was collected through the 10 objective (Mitutoyo, Kanagawa, Japan), passed through a cross-polariser (LPNIRE200-B, Thorlabs) mounted on a rotation stage (LCRM2/M, Thorlabs), relayed via an infinity-corrected tube lens (TTL200-B, Thorlabs), and imaged onto a CMOS camera (Basler aca2040-90um NIR, 5.5 x 5.5 µm pixels, Basler, Ahrensburg, Germany). Images (1024 512 pixels) were acquired with a frame rate of 194 frames/s and an exposure time of 5000 µs. Each acquisition lasted 120 s and was performed from month 13 to month 15.

Spatial speckle contrast was calculated for each frame using a 5 × 5 pixel sliding window and converted to BFI values^23^. Representative cardiac cycles were extracted from the temporal contrast signal by frequency-based cycle identification, followed by quality-control filtering and temporal interpolation to generate an averaged cardiac cycle for each recording (Fig 2c).

PI maps were subsequently calculated from BFI signals as:

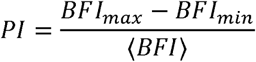

Vascular segmentation was performed manually using reference images obtained from the surgical microscope together with bolus arrival time and intravascular oxygen pressure measurements derived from 2P recordings (see below). Arterial, venous, and parenchymal regions of interest to quantify mean BFI, BFI-derived PI, and RI.

All preprocessing and analysis procedures were implemented using a custom analysis pipeline described previously^7^.

#### Two-photon imaging

2P was performed with a scope Prairie Ultima-IV (Bruker Inc., Madison, WI, USA), equipped with a 4-channel photomultiplier detector system. We used an Olympus 25x water immersion objective (1.05 NA; 2 mm working distance). Fluorescence dyes were excited with a tunable Ti: Sapphire laser, Chameleon Ultra II (Coherent, Santa Clara, CA, USA). For this study, excitation wavelengths of 900 nm and 930 nm were used to excite the fluorescent dyes Texas Red dextran (70 kDa; D1864, Thermo Fisher Scientific, Waltham, MA, USA) and the oxygen probe Oxyphor2P (Oxy2P), respectively, as described below. The scope was driven by Prairie View Software version 5.1 (Bruker).

To extend the depth of focus and reduce sensitivity to axial motion inherent to scanning techniques, thus to 2P with our current objective, we incorporated a home built Bessel beam module into our microscope, adapted from a previously validated design²³. An axicon (AX121-B) generates an annular beam profile, which is relayed through two achromatic doublet lenses (AC254-125-B-ML and AC254-500-B-ML) before entering the scope scanning head, forming an annular illumination pattern at the back aperture of the objective. The properties and positions of axicon and the relay lenses are calculated to match the original, Gaussian, beam diameter at the back aperture of the objective. Upon focusing by the objective, annular beam profile is transformed into Bessel beam with extended depth of focus. While scanning, the extended depth of focus provides the volume image in one scan, with deliberately destroyed axial, but with preserved lateral resolution. This configuration is not sensitive to axial vessel movements across the imaging volume and improved the fidelity of dye-dilution curves and vessel diameter estimates, both of which are degraded in conventional Gaussian-beam 2P due to axial out-of-focus displacement of the vessels.

Before detection, the emission signal was reflected using an 875-nm long-pass dichroic and filtered with an 875-nm blocking filter. A 690-nm dichroic separated far-red signal from red-shifted signal. Emission from each probe was further isolated using a 605/70 bandpass filter for Texas Red and a 760/50 bandpass filter for Oxy2P (all Chroma Technology Corp, Bellows Falls, VT, USA). Both signals were detected with GaAsP-PMTs: H10770PB-40 for Texas Red and H10770PB-50 for Oxy2P (Hamamatsu, Hamamatsu City, Japan). Each probe was administered via a PE-10 catheter placed in the mouse’s tail and secured with dental cement (GC Fuji PLUS; GC Corporation, Tokyo, Japan), as previously described^13^.

#### Intravascular oxygen pressure

For intravascular pO2 estimates, phosphorescent probe Oxy2P^24^ (5 mM final concentration in the blood plasma; ≈50 µl of 200 mM stock solution per mouse) was injected via the tail catheter to ensure sufficient signal. Ten points were placed within pial vessels (one per vessel) in the Phosphorescence PointScan mode of the acquisition software. Single points were excited at 960 nm using a 10 µs-long gate, and phosphorescence was collected over a 300-µs window, with the initial 10 µs (coincident with excitation) excluded, yielding 290 µs of usable signal. Each excitation and collection sequence, consisting of the excitation gate followed by the 300 µs phosphorescence window, was repeated 55 times per point, and this block was run for 300 iterations. This gave 16,500 excitation cycles per point (55 x 300). The photons detected after each cycle were accumulated into a single decay histogram per point by time-correlated single-photon counting, and the phosphorescence lifetime was estimated from this summed decay curve. Phosphorescence emission was detected using a GaAsP-PMT. The PMT output was channeled via amplifier/discriminator to a photon counting card (SPC-150; Becker & Hickl GmbH, Berlin, Germany), as previously described^24^. Intravascular point scans (i.v. pO2) were processed using a custom-made MATLAB GUI (tpmpO2)^13^ to calculate pO2 values (mmHg) from individual points inside vessels in the field of view (resolution, 512 × 512 pixels). Conversion of phosphorescence decay times to pO2 values was based on calibrations of Oxyphor 2P performed in independent experiments.

#### Vessel diameter

Time-series scans were acquired at 10 frames/s using the spiral-scan mode of PrairieView software, over a field of view of 256 × 256 pixels (589 µm²), with an estimated x-y resolution of 2.65 µm and an estimated axial resolution of 0.88 µm (refractive index = 1.4). Each scan lasted 25 s, and two scans were performed per mouse. These scans were originally designed to assess functional activation; here, only the 5 s pre-stimulus baseline period was analyzed, as the present study focused on steady-state measurements.

For analysis, vessels were manually identified and classified as pial arteries and veins, diving arteries and draining veins, based on whether they projected into deeper tissue or ran parallel to the cortical surface. Arteries and veins were distinguished based on bolus arrival time (earlier arrival in arteries; see Indicator-dilution) and intravascular pO₂. The same images used for pO_2_ and indicator-dilution measurements were used for diameter estimation.

Vessel diameter was estimated using FIJI (National Institutes of Health, Bethesda, Maryland, USA) and the VasoMetrics macro^25^. Before diameter estimation, signal-to-noise ratio and motion were corrected using contrast enhancement and rigid image registration, integrated into a custom-made macro previously described^13^. The user drew a through-line along the axis of the vessel of interest, from which VasoMetrics generated perpendicular cross-lines^25^. Cross-line length was set manually as the vessel diameter (in pixels) plus an additional 2 to 5 pixels to account for changes; the spacing between cross-lines was set manually to 1 to 3 pixels, based on the length of the through-line.

VasoMetrics measures vessel diameter from the fluorescence intensity along each cross-line, generating an intensity profile normalized between zero and one (obtained by subtracting the minimum intensity and dividing by the maximum). Vessel diameter was calculated as the full-width at half-maximum (FWHM): the distance, in pixels, between the points where the intensity profile crosses 50% of its maximum on the rising and falling edges, corresponding to the lumen boundaries. This was applied to each cross-line in every frame of the time series. VasoMetrics outputs the diameter for each cross-line, as well as the mean diameter across all cross-lines with the associated standard deviation (SD).

#### Indicator-dilution

An indicator-dilution technique was applied to estimate MTT and CTH as previously described^13,26^. For this study, we acquired time series at 10 10 frames/s over a FOV of 256 x 256 pixels (2.65 µm x-y pixel size; 0.88 µm z-size). Boluses of 10 µl of Texas Red (5 mg/ml in saline) were injected through the catheter 7 s into the time-series scan to establish a pre-bolus baseline before dye arrival.

MTT and CTH were estimated by modeling the vascular transport function (TF) through deconvolution of the dye bolus passage between a manually selected arterial input function (AIF) and venous output function (VOF), as previously described^13,26^. The TF was modeled as a gamma probability density function, with MTT defined as the mean (α·β) and CTH as the standard deviation (√α·β). Coefficient of variance (COV = CTH/MTT) was estimated as a normalized marker of transit time heterogeneity.

#### Ex-vivo MRI scans

Four mice per group were selected for in-skull ex vivo MRI. Following euthanasia, mice were perfusion-fixed with heparinized saline (10 U/mL) followed by 4% paraformaldehyde (PFA). Heads were then prepared for in-skull imaging by removing soft tissue and the lower jaw, followed by post-fixation for weeks. Prior to imaging, samples were washed in phosphate-buffered saline (PBS) to improve signal by removing excess fixative and transferred to an imaging tube filled with fluorinated liquid (FC-770; 3M) to minimize image susceptibility artifacts.

MRI data were acquired using a 9.4 T Bruker BioSpec system equipped with a 15 mm volume coil. High-resolution structural images were obtained using a segmented T2-weighted RARE sequence with an in-plane resolution of 150 µm and a slice thickness of 200 µm. Imaging parameters included repetition time (TR) = 4500 ms, echo time (TE) = 31.7 ms, and 16 signal averages.

Brain regions, including somatosensory cortex (SMS), hippocampus (HC), white matter (WM), caudate-putamen, and thalamus, were manually segmented using ITK-SNAP software, as previously described^27^. Regional volumes were calculated by multiplying the number of segmented voxels by the nominal voxel volume (150 x 150 x 200 µm), normalized by the total brain volume.

#### Spatial Transcriptomics and MOFA-MEFISTO analysis

For spatial transcriptomics and subsequent immunohistochemistry, mice were sacrificed by an overdose of pentobarbital following induction of anesthesia with 5% isoflurane. The brain was dissected and harvested, and the left hemisphere was mounted in OCT compound for cryosectioning. Coronal sections were cut at 10 µm thickness, approximately −8 mm from the olfactory bulb, and mounted on Visium Spatial Gene Expression slides (10x Genomics). Sections were fixed in methanol, stained with hematoxylin and eosin (H&E), and imaged at 20x magnification.

*Visum library generation:* Spatial gene expression libraries were generated with the Visium Spatial Gene Expression kit (10x Genomics). One left-hemisphere cryosection per animal was placed within a 6.5 × 6.5 mm capture area. Permeabilization time was first optimized with the Visium Spatial Tissue Optimization kit (10x Genomics), testing 3, 6, 12, 18, 24, and 30 min alongside positive and negative controls. An 18-min permeabilization was selected.

Sections were methanol-fixed, H&E-stained according to the 10x Genomics protocol (CG000160, Rev B), and imaged using an Olympus VS120 slide scanner. Visium Spatial Gene Expression libraries were prepared according to the manufacturer’s protocol (CG000239, Rev D). Tissue was permeabilized for 18 min, followed by on-slide mRNA capture, reverse transcription, and second-strand synthesis. cDNA amplification cycles were determined by qPCR, and cDNA was amplified for 17 cycles. Libraries were generated from 10 µL of amplified cDNA by fragmentation, end repair, A-tailing, adaptor ligation, and 13 cycles of sample-index PCR using unique dual indexes (Dual Index Kit TT Set A, 10x Genomics). Final libraries were quantified using a Qubit fluorometer (Thermo Fisher Scientific), and fragment size distributions were assessed using a Bioanalyzer (Agilent). Libraries were pooled in equimolar ratios and sequenced on a DNBSEQ-G400 platform (BGI). Sequencing was paired-end, 100 + 100 bp with 10 + 10 bp dual index reads.

*Read alignment and quantification*: FASTQ files were processed using Space Ranger v2.1.1 (10x Genomics) with the Visium v1 3′ chemistry and mm10 reference (refdata-gex-mm10-2020-A). Fiducial alignment and tissue-covered spots were manually defined in Loupe Browser. Reads were aligned using STAR, assigned to spatial barcodes, and UMIs collapsed to generate gene-by-spot count matrices.

*Data processing and region annotation:* Downstream analyses were performed in R (v4.2.3) with Seurat (v4.3.0). Filtered feature–barcode matrices and H&E images were loaded with Load10X_Spatial, and all spots under tissue were retained. The count matrices were merged and normalized with SCTransform (sctransform v0.3.5). Principal component analysis was performed, and the first 20 PCs were used to build a shared nearest-neighbor graph, followed by Louvain clustering (FindClusters) and UMAP embedding.

Anatomical regions were defined from the unsupervised clusters. The cluster assignments were then refined manually in Loupe Browser, using the H&E images and the Allen Mouse Brain Atlas as reference. Twelve regions were annotated: upper cortical layers (CTX_tr1-3), deep cortical layers (CTX_tr4-6), the most superficial cortical/pial region (PIA), hippocampal formation (HPF), amygdala (AMG), piriform area (PIR), caudoputamen (PUT), thalamus (TH), hypothalamus (HYP), medial habenula (MHb), white matter (WM), and lateral ventricle (LV).

*Region-specific differential expression analysis: S*ex-specific differential expression was assessed within each annotated region using Seurat FindMarkers on SCT-normalized data with a two-sided Wilcoxon rank-sum test. All 22,611 detected genes were tested without prefiltering, and P values were Bonferroni-adjusted within each region. Genes with adjusted P < 0.05 were considered differentially expressed; positive and negative log2 fold changes indicated female– and male-enriched expression, respectively.

*Gene ontology enrichment analysis:* Female– and male-enriched DEG sets from each region were tested for over-representation of Gene Ontology Biological Process terms. The test used enrichGO in clusterProfiler (v4.6.2) with the org.Mm.eg.db annotation (v3.16.0) and gene symbols as identifiers. Enrichment was tested only for gene sets with more than 10 genes, against a background of all GO-annotated genes in org.Mm.eg.db. P values were adjusted with the Benjamini–Hochberg method, and terms with adjusted P < 0.05 were considered significantly enriched.

*Method for Functional Integration of Spatial and Temporal Omics (MEFISTO):* Spatial transcriptomic data were analyzed from the processed Seurat object. Normalized SCT expression values were extracted and paired with spot-level spatial coordinates from the image metadata. MEFISTO input matrices were constructed with genes as features and spots as observations. The four tissue sections were modeled as separate groups. Spatial image coordinates were scaled within each sample and supplied as two continuous covariates to model spatially structured variation. A Gaussian likelihood was used for the normalized expression values. The model was initialized with 10 latent factors and trained with MEFISTO to identify spatially structured transcriptional programs across spots. Five factors were retained after training. Because Factor 5 was less clearly interpretable, downstream visualization focused on Factors 1-4. For each factor, factor values were mapped back onto spatial coordinates, summarized across annotated anatomical regions, and compared between females and males within the regions showing the highest positive mean factor values. Gene weights were extracted to identify genes contributing most strongly to the positive and negative ends of each factor. Spot-level female-versus-male differences were assessed within selected regions using Wilcoxon tests and are reported as descriptive statistics.

#### ELISA

Blood was collected into tubes pre-rinsed with heparinized saline (50 U/ml), centrifuged at 1,500 × g for 10 min at 4 °C, and plasma was aliquoted and stored at −80 °C until analysis. Concentrations were measured with commercial kits according to the manufacturers’ instructions. Estradiol was measured with the QuicKey Pro Mouse E2 ELISA kit (Elabscience; undiluted). Corticosterone was measured with the DetectX Corticosterone ELISA kit (Arbor Assays, 1/100 dilution). MMP9 was measured with the Quantikine ELISA kit (Biotechne, R&D Systems; 1/20 dilution). Concentrations were interpolated from four-parameter logistic standard curves.

#### Immunohistochemistry

Immunohistochemistry was performed on tissue from two sources: parallel sections from the four mice also used for spatial transcriptomics (2 females, 2 males), and additional mice that were aged to 15 months but not included in the ex vivo MRI cohort (3 females, 2 males), yielding a total sample size of N = 5 females and N = 4 males.

Slides were air-dried for 2 h at 37 °C and rinsed in PBS (3 × 5 min). Antigen retrieval was performed in Tris/EDTA buffer (10 mM Tris base, 1 mM EDTA, 0.05% Tween-20, pH 9.0) at 70 °C for 10 min, followed by 20 min at room temperature in the same solution. Sections were marked with a hydrophobic barrier pen, and free aldehydes were quenched with 50 mM NH₄Cl in PBS for 10 min. Sections were rinsed in PBS-T (0.1% Triton X-100 in PBS) and blocked for 60 min at room temperature in blocking buffer (1% milk, 3% BSA in 0.1% PBS-T). Sections were incubated overnight at 4 °C with primary antibodies diluted in 1% BSA in PBS (see below), rinsed in PBS-T (3 × 5–10 min), and incubated with secondary antibodies diluted in 1% BSA in PBS (see below) for 2 h at room temperature. Sections were rinsed in PBS with DAPI (300 nM) for 10 min, rinsed in PBS for 5 min, treated with TrueBlack® lipofuscin autofluorescence quencher (Fremont, CA, USA; 1:20 in 70% ethanol) for 1 min to reduce background autofluorescence, rinsed again in PBS (10 min) and distilled water, and mounted using Dako Fluorescent Mounting Medium (Agilent Technologies, Santa Clara, CA, USA) with coverslips sealed with nail polish.

For vascular and astrocytic labeling, sections were co-stained with goat anti-CD31 (R&D Systems, AF3628; 1:200) and chicken anti-GFAP (Aves Labs, GFAP, RRID: AB_2313547; 1:500), followed by donkey anti-goat IgG Alexa Fluor 568 (Invitrogen, A-11057; 1:1000) and donkey anti-chicken IgY Alexa Fluor 488 (Invitrogen, A78948; 1:500). For microglial labeling, sections were co-stained with goat anti-CD31 (R&D Systems, AF3628; 1:200) and rabbit anti-Iba1 (Abcam, ab178847; 1:500), followed by donkey anti-goat IgG Alexa Fluor 568 (Invitrogen, A-11057; 1:1000) and goat anti-rabbit IgG Alexa Fluor 488 (Invitrogen, A32731; 1:1000).

#### Barnes maze

We applied the Barnes maze to assess visuospatial learning and memory in mice. Our protocol was based on previous studies that used both bright light and a white-noise stimulus to motivate escape from the open platform to a small dark chamber, a brightly lit maze associated with aversive noise^28–30^. The test comprises a raised light-blue antiglare platform (ø = 1 meter), with a textured surface, positioned 91.5 cm above the ground. The platform had twenty 5-cm holes evenly spaced around the edge, located 2.5 cm from the perimeter. During training sessions, a dark plastic escape box (10×8.5×4 cm) was placed under one of these holes, and its position remained consistent throughout trials. Visual cues were positioned in the testing environment and remained constant across the testing period. Illumination was provided by an overhead lamp (600 lux at the surface of the maze) positioned above the center of the maze. The total trial time spanned 6 consecutive days: one habituation day on day 0, 9 training trials from day 1 to day 3 (3 trials per day), a resting day on day 4, and the final probe on day 5 (Fig. 1a). During the habituation phase, mice were introduced to the maze for 3 min, then allowed to explore the dark plastic escape box for 2 min. Training days featured three trials; the animal was placed in the center of the maze in a plastic start box (15 cm in diameter). After 10-15 s, the box was lifted, and the animal had 3 min to locate and enter the designated escape hole. Upon successful entry, the mouse was left in the escape box for 1 min before being moved to the home cage. If the mouse failed to find the escape hole within the allotted time, it was gently guided to the target hole beneath the inverted beaker and left there for 1 min. On day 5, a three-minute probe trial was conducted like the training trials, except the escape box was removed.

**Figure 1.**
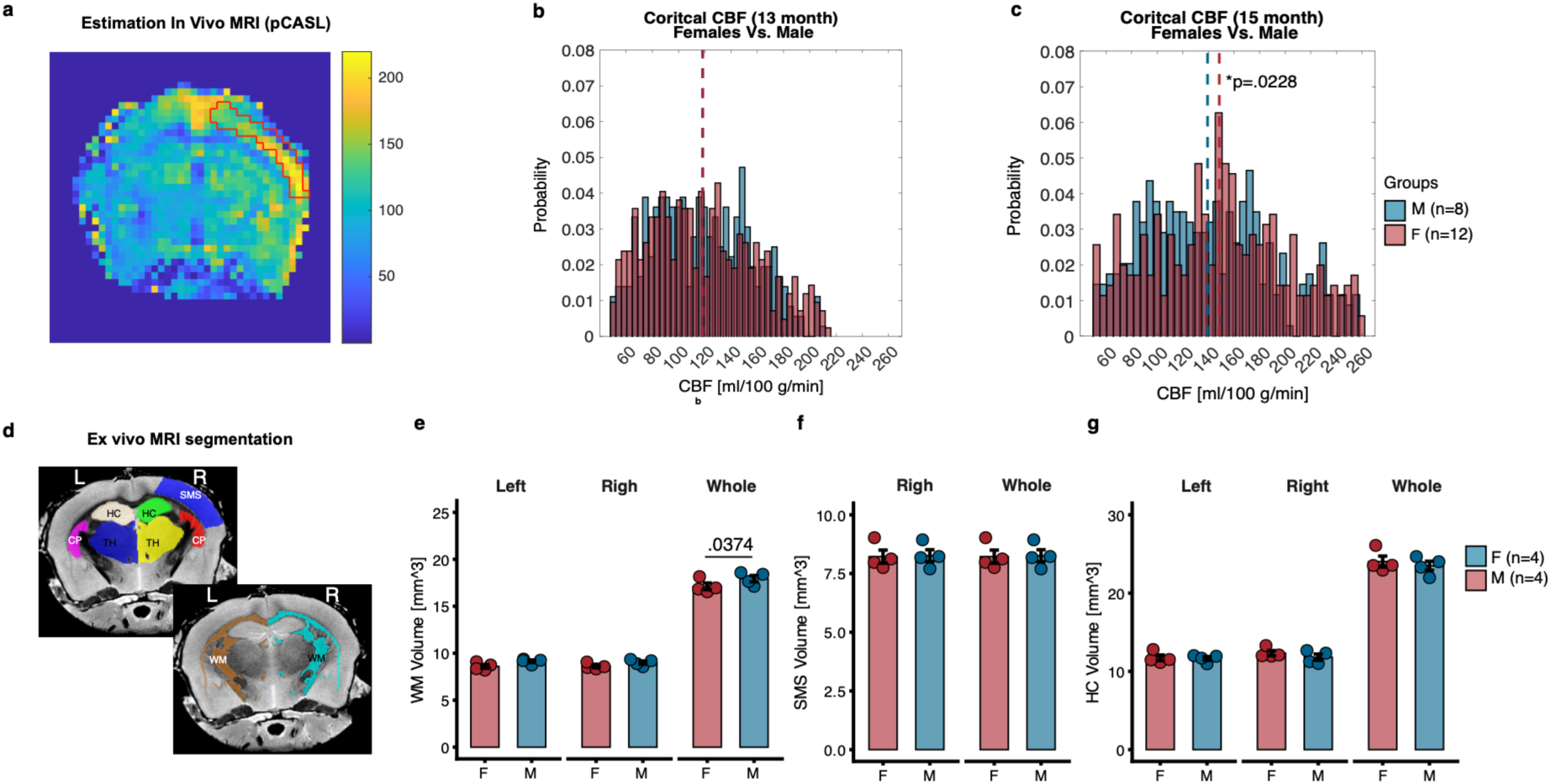
Sex-dependent changes in CBF and brain volumetrics. **(a)** Example of CBF estimate from pCASL sequence in a central brain slice. Cortical CBF in female and male mice at 13ᵗʰ **(b)** and 15ᵗʰ months. **(c)** Permutation tests (100,000 permutations) with t-tests of means: M13, F vs. M, n.s. (p = 0.8951); M15, F vs. M diff. = 9.18 ± 3.94 ml/100 g/min, p = 0.0228. Distributions show pooled CBF values (pixel-wise) from all subjects, color-coded by group. **d.** Regional volumetrics based on ex vivo MRI. Bilateral segmentation of hippocampus (HC), caudoputamen (CP), thalamus (TH), and white matter (WM) was conducted. The right somatosensory area (SMS) was also segmented. Comparisons in **e.** WM, LMM: WM whole brain F vs. M diff. = – 0.8181 ± 0.345 mm^3^, d = –2.577; **f.** SMS, and **g.** HC. Error bars: mean ± SEM. F: female, M: male. Statistical results are provided in the supplementary material.

Behavior was recorded via an overhead camera for subsequent analysis (Ethovision XT 11.5 software, Noldus, Wageningen, the Netherlands). Key metrics, including errors, latency (to first goal visit), distance traveled, velocity, time immobile, and time spent in entry zone, were measured as indicators of spatial memory. Barnes maze was performed before habituation for awake imaging at months 13, 14, and 15.

#### Statistics

All statistical analyses were performed with R version 4.5.1. A table of statistical analyses for each figure panel is provided in the supplementary materials.

CBF group differences were analyzed using nonparametric permutation tests. Within-group age comparisons (13 vs. 15 months) were performed separately for males and females, and between-group sex comparisons (males vs. females) were performed at each age. All tests were conducted on pooled data from all subjects in each group, using 100,000 permutations to estimate empirical p-values as previously applied^31^. Because only one perfusion parameter was tested, p-values are reported uncorrected.

For in vivo imaging in LSCI-derived BFI, PI, RI, and HR, and in pial vessel diameter and its coefficient of variation (COV), were each tested using LMM, with the respective parameter modeled as a function of sex, month, and their interaction, and animal ID included as a random intercept to account for repeated measurements across time points (e.g., pi ∼ sex * months + (1|ID)). Separate models were fitted for arteries, veins, and parenchymal regions. Sex differences in MTT, CTH, and COV were each tested separately using LMM, with the respective parameter modeled as a function of sex, and animal ID, artery, and vein included as random intercepts to account for repeated measurements within mice and variability across individual vessels (e.g., mtt ∼ sex + (1|MouseID) + (1|artery) + (1|vein)). The relationship between CTH and MTT was analyzed using a separate LMM, with CTH modeled as a function of MTT, sex, and their interaction, and animal ID included as a random intercept (cth ∼ mtt * sex + (1|ID)). To assess the robustness of the MTT × sex interaction, a leave-one-out sensitivity analysis was performed by refitting the model after sequential exclusion of each animal and tracking the resulting interaction estimate. Sex differences in intravascular pO2 and OEF were tested using LMM, with each parameter modeled as a function of sex, vessel segment identity (e.g., pial or diving artery/vein), and their interaction, and animal ID included as a random intercept (e.g., po2 ∼ sex * vessel_segment_ID + (1|MouseID)).

All models were fit using maximum likelihood estimation. Residual diagnostics, including residual-versus-fitted plots and Q-Q plots of residuals, were visually inspected to verify model assumptions. Estimated marginal means and pairwise sex contrasts were computed using the emmeans package, performed both across all months and within each month separately where applicable; within-sex comparisons across months were additionally conducted to assess longitudinal changes, with BH correction applied for multiple comparisons in these longitudinal within-sex analyses. Cohen’s d was calculated as the model-estimated contrast divided by the residual standard deviation of the fitted model.

Sex differences in immunohistochemical markers (CD31 coverage, GFAP area fraction and intensity, Iba1 area fraction and intensity) and in ex vivo MRI regional brain volumes were assessed using a permutation test. For each comparison, the test statistic was the absolute difference between group means; a null distribution was generated by randomly shuffling group labels and recomputing this statistic over 10,000 permutations, and the p-value was calculated as the proportion of permuted differences equal to or exceeding the observed difference. Effect sizes were calculated as Cohen’s d.

Sex differences in plasma corticosterone, CXCL13, MMP-9, IgA, and IgG concentrations were assessed using the Wilcoxon rank-sum test.

Barnes maze acquisition across training sessions was analyzed using linear mixed-effects modeling. Error counts were modeled as a function of sex, training session, month, and their interactions. The training session was treated as a continuous numeric variable to evaluate the learning rate across sessions. The model included animal ID as a random effect, with both random intercepts and random slopes for session to account for individual differences in baseline performance and learning trajectory (e.g., errors ∼ sex * session_num * month + (1 + session_num | ID).

Learning within each sex and month was assessed using estimated marginal trends from the fitted model. Session slopes were extracted with emtrends, providing the change in errors across training sessions for each sex-by-month group. Negative slopes indicate a reduction in errors over training and were interpreted as evidence of learning. Sex differences in learning rate were tested by comparing session slopes between males and females within each month. Pairwise comparisons were adjusted using the BH correction. Cohen’s *d* was estimated using the estimated slope or slope difference divided by the residual standard deviation of the model.

Barnes maze probe day performance was analyzed using linear mixed-effects models implemented in the lme4 package in R. The number of errors was modeled as a function of sex, month, and their interaction, with animal ID included as a random intercept to account for repeated measurements across animals (e.g., errors ∼ sex * month + (1|ID)). Models were fit using maximum likelihood estimation. Residual diagnostics, including residual versus fitted value plots and Q–Q plots, were visually inspected to confirm model assumptions.

Estimated marginal means were calculated using the emmeans package. Pairwise post hoc comparisons between sexes were performed both across all months and within each month separately. Benjamini–Hochberg (BH) correction was applied to control for multiple comparisons. Effect sizes were estimated using Cohen’s *d*, calculated from the model-estimated contrasts divided by the residual standard deviation of the fitted model.

Associations between Barnes maze probe day error counts and behavioral performance measures were assessed using Pearson correlation analyses. Correlations were performed separately for each sex and month to evaluate whether relationships differed across groups and time points.

Two correlation analyses were conducted: errors versus time immobile, and errors versus time spent in the goal quadrant. For each sex-by-month subgroup, animals with missing values for either variable were excluded. Correlations were calculated only when at least 3 animals were available in a subgroup. Pearson’s correlation coefficient (*r*) and corresponding *p*-values were extracted using cor.test in R.To account for multiple testing across sex and month subgroups, *p*-values were adjusted using the Benjamini–Hochberg false discovery rate correction.

All LMM were fit using maximum likelihood estimation. Residual diagnostics, including residual-versus-fitted plots and Q–Q plots of residuals, were visually inspected to verify model assumptions.

Data are presented as mean ± SEM. Statistical significance was defined as *p* < 0.05.

## Results

### Cortical CBF is increased in female mice at 15 months old

We first asked if there were any sex-dependent differences in cerebral hemodynamics. We therefore performed in vivo MRI and estimated absolute CBF in the mouse cortex using an established protocol in our lab for pseudo-continuous arterial spin labeling (pCASL)^20^. CBF was estimated only in the hemisphere opposite the implanted cranial window (right hemisphere – Fig. 1a).

We did not observe sex differences in cortical CBF at 13 months (Fig. 1b; female: 118.53 ± 40.89 ml/100 g/min; male: 118.14 ± 37.05 ml/100 g/min; p = 0.8951). However, by 15 months, female mice exhibited significantly higher CBF compared with males (Fig. 1c; female: 148.49 ± 52.68; male: 139.31 ± 49.65; p = 0.0228), indicating a sex-dependent increase in cortical perfusion with aging.

Pixel-wise analysis of cortical CBF revealed significant increases from 13 to 15 months in both females and males (Fig. S1). In females, mean CBF rose by ∼25.3%, while in males it increased by ∼18%.

### Sex differences in cerebrovascular pulsatility diminish with aging

Having observed increased cortical CBF in females at 15 months old, we next asked whether this reflected altered vascular tone. Previous studies have shown that changes in pulsatility index (PI) contribute to brain pathology and the development of dementia^32,33^. We, therefore, performed laser speckle contrast imaging (LSCI) to evaluate cerebrovascular dynamics by quantifying blood flow index (BFI) and deriving pulsatility-related parameters across arterial, parenchymal, and venous regions (Fig. 2a-c).

**Figure 2.**
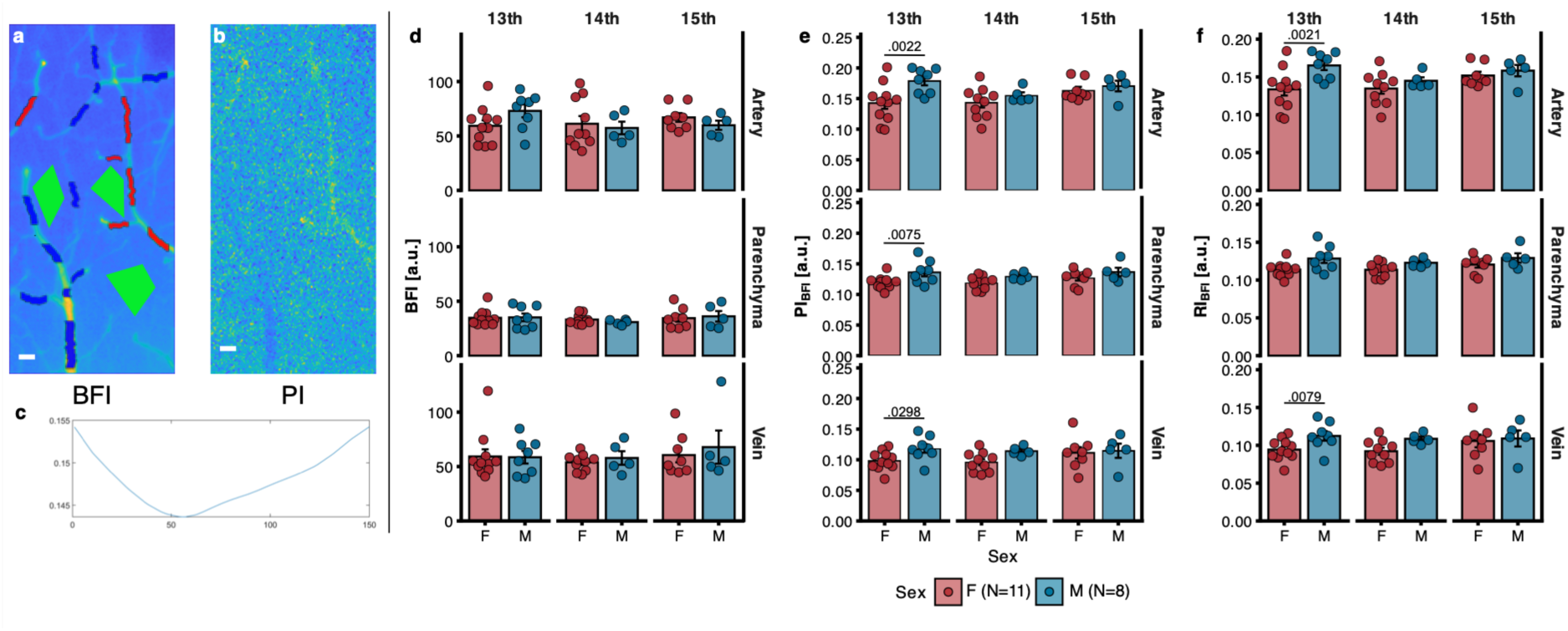
Sex– and age-dependent changes in blood flow index (BFI) and pulsatility index (PI) using LSCI. **a**. Representative average BFI image with arterial (red), parenchyma (green), and vein (blue) ROIs. **b.** Representative average PI image. **c.** Average contrast cycle over the analyzed frames. **d.** Average BFI per mouse and region. **e.** Average PI per mouse and per region. **f.** Average PI per mouse and per region. Statistical analysis: LMM; post hoc comparisons by emmeans with BH correction, assessing effects per region. F, female; M, male; bpm, beats per minute. Error bars: mean ± s.e.m. Statistical results are provided in the supplementary materials.

We did not observed sex differences in BFI across vascular compartments or ages (Fig. 2d), and no consistent age-dependent changes were detected within either sex, indicating that baseline cortical perfusion remained relatively stable between 13 and 15 months.

In contrast, analysis of the pulsatility index derived from BFI^7^ revealed clear sex differences at 13 months (Fig. 2e). Male mice exhibited significantly higher PI compared with females in arteries (p = 0.0022), parenchyma (p = 0.0075), and veins (p = 0.0298). No significant sex differences were detected at 14 or 15 months. These findings extend our previous observations in 12-month-old mice, in which females exhibited lower microvascular PI than males despite comparable perfusion^7^. In the present cohort, this difference is most evident at 13 months and diminishes by 15 months of age, reflecting a slight decrease in males from 13 to 14 months (p = 0.0689).

A similar pattern was observed for the BFI-derived resistance index (Fig. 2f), with males showing significantly higher values than females at 13 months in arteries (p = 0.0021) and veins (p = 0.0079), whereas no sex differences were detected at later ages. Estimations of heart rate derived from the LSCI pulsation signal did not differ significantly between sexes or across ages (Fig. S2), indicating that the observed differences in pulsatility were not attributable to systemic changes in cardiac rhythm.

### Vessel diameter does not differ between sexes at 15 months

Having found that sex differences in PI converge by 15 months, we next asked whether this convergence extended to vessel structure on the superficial layers of the brain. Using two-photon imaging (2P), we examined pial and diving vessel diameters and short-term diameter variability in arteries and veins. A total of 55 arterial and 65 venous segments were scanned twice and analyzed across 13 mice (female N = 9; male N = 5). Vessels were classified according to their anatomical relationship with the cortex as pial or diving, with pial vessels defined as those running parallel to the cortical surface and diving vessels as branches entering or exiting deeper cortical layers (Fig. 3a).

**Figure 3.**
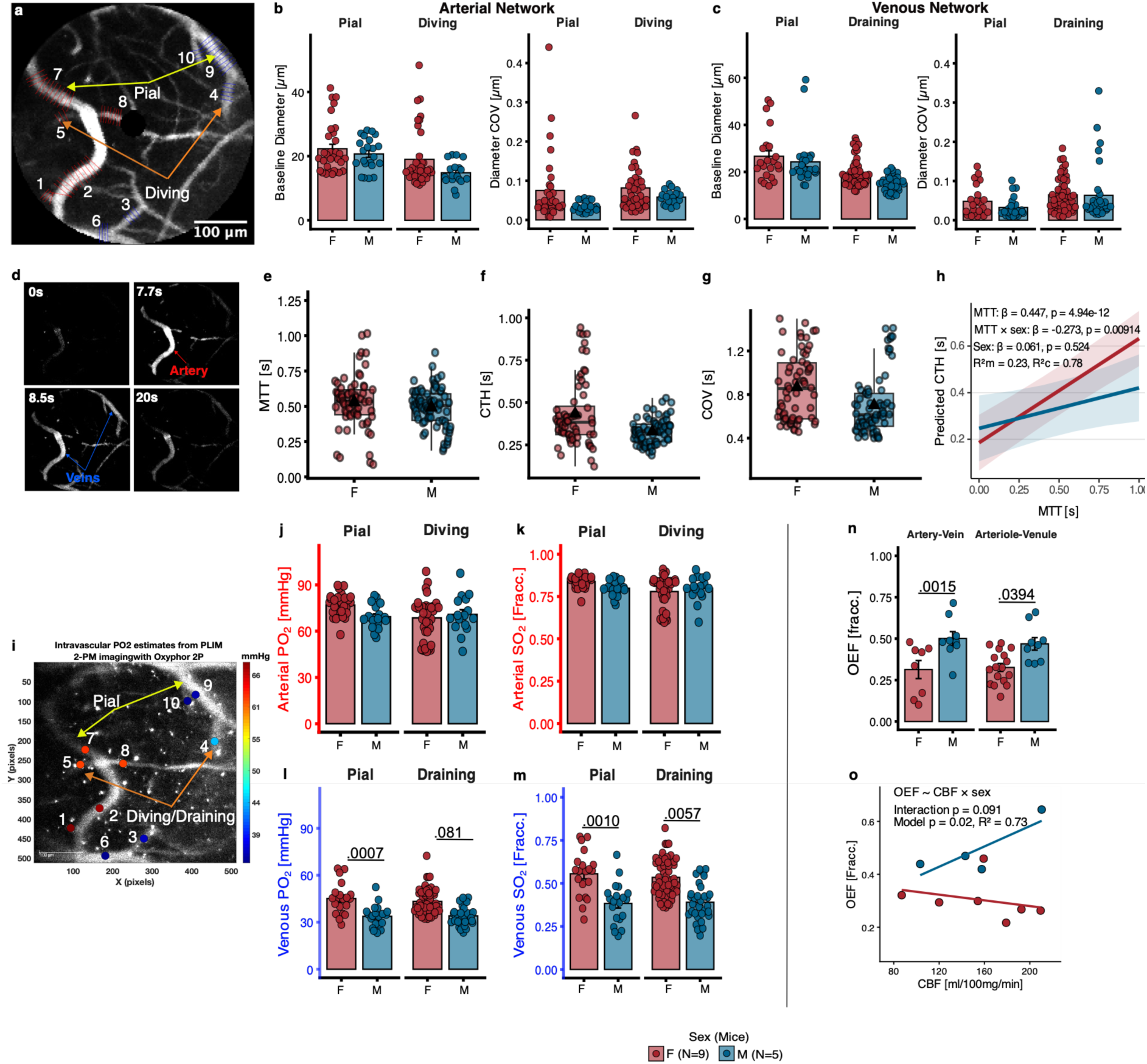
Sex differences in cortical oxygen extraction and capillary transit dynamics despite similar vessel diameters. **a**. Representative 2P image of cortical vasculature showing vessels used for diameter, oxygen measurements, and estimation of capillary transit times. **b–c.** Baseline vessel diameter and diameter coefficient of variation (COV) measured in pial and diving arterial segments **(b)** and pial and draining venous segments **(c)** using the ImageJ macro VasoMetrics from 5-s 2P time series. Points represent individual vessels; bars indicate mean ± SEM. No significant sex differences were detected. **(d)** Representative frames illustrating the propagation of a fluorescent bolus between an arterial input and venous output during indicator-dilution imaging. **(e–h)** Capillary transit dynamics derived from the indicator-dilution model. MTT **(e)**, CTH **(f)**, and COV **(g)** are shown for individual vessels across mice. **(h)** Relationship between MTT and predicted CTH derived from linear mixed-effects modeling. A significant MTT × sex interaction (*p* = 0.009) indicates sex-dependent coupling between MTT and CTH. **i.** Example intravascular PO₂ map obtained using 2-PLIM after administration of the oxygen-sensitive dye Oxyphor2P**. j–k** Arterial intravascular oxygenation. Arterial PO₂ **(j)** and calculated SO₂ **(k)** were estimated from phosphorescence lifetime decay fits in pial and diving arteries. No significant sex differences were observed. **l–m.** Venous oxygenation. Venous PO₂ **(l)** and SO₂ **(m)** were significantly higher in females compared with males in both pial and draining veins (LMM). **n.** Estimation of OEF from paired arterial and venous networks per mouse showing lower OEF levels in females compared with males in both pial and diving/draining regions. **o.** Relationship between cortical cerebral blood flow (CBF) measured by MRI and OEF derived from 2P imaging. The relationship differed between sexes (model: OEF ∼ CBF × sex, interaction *p* = 0.091), indicating sex-dependent coupling between perfusion and oxygen extraction. Error bars: mean ± SEM. F: female, M: male, MTT: mean transit time, CTH: capillary transit-time heterogeneity, COV: coefficient of variance. Statistical results are provided in the supplementary material.

During the 5-s time series, arterial baseline diameter did not differ significantly between sexes in either pial or diving arteries (Fig. 3b). Similarly, the coefficient of variation (COV) of arterial diameter, reflecting short-term diameter fluctuations during the acquisition window, did not differ between sexes for either vessel type. Analysis of the venous network yielded comparable results. Baseline venous diameter and COV did not differ between sexes in pial or diving veins (Fig. 3c). These findings indicate that vessel structure and short-term caliber fluctuations do not account for the observed differences in cerebral hemodynamics, motivating an examination of capillary flow dynamics.

### Capillary flow heterogeneity couples more steeply with transit time in female microvascular networks

Having found no sex differences in pial vessel diameter or short-term caliber variability, we next asked whether capillary flow dynamics differed by sex. Using an indicator-dilution framework developed in our laboratory, we quantified cerebral microvascular transit dynamics as mean transit time (MTT) and capillary transit-time heterogeneity (CTH) between arteries and veins at steady state (Fig. 3d)^13,26,34^.

At steady state, microvascular flow parameters were comparable between sexes, with no significant differences in MTT, CTH, or COV between female and male mice (Fig. 3e–g). These findings indicate that, at steady state, the overall distribution of capillary transit times does not differ significantly between female and male mice.

Across all measurements, CTH increased strongly with MTT (β = 0.447, *p* = 4.94 × 10⁻¹²), demonstrating that longer capillary transit times are associated with greater perfusion heterogeneity (Fig. 3h). Importantly, the relationship between MTT and CTH differed significantly between sexes (MTT × sex interaction: β = −0.273, *p* = 0.009), indicating sex-dependent coupling between transit time and microvascular flow heterogeneity. Specifically, the slope relating CTH to MTT was steeper in females than in males, suggesting that increases in MTT are associated with a stronger rise in CTH in female microvascular networks. The model explained a substantial fraction of the variability in CTH (marginal R² = 0.23; conditional R² = 0.78).

This steep, near-proportional coupling in females is consistent with a passive microvascular network, in which CTH tracks MTT directly rather than being actively constrained, as predicted by models of capillary flow homogenization and oxygen extraction efficiency^9^. By contrast, the shallower slope in males suggests flow regulation that limits the rise in CTH as MTT increases, even under steady-state conditions. Collectively, these findings indicate that although average steady-state MTT, CTH, and COV do not differ between sexes, female microvascular networks exhibit a disproportionate rise in flow heterogeneity as transit time increases, potentially reflecting reduced active flow regulation. This pattern could compromise oxygen-extraction efficiency and contribute to the elevated cortical CBF observed in females at 15 months.

### Higher venous PO_2_ in females drives reduced OEF

Having found that female microvascular networks exhibit a steeper coupling between CTH and MTT, we next asked whether this altered flow organization was accompanied by differences in intravascular oxygenation and oxygen extraction fraction (OEF). We therefore measured the intravascular oxygen partial pressure (PO_2_) using two-photon phosphorescence lifetime microscopy (2PLM)^35^ after infusion of the oxygen-sensitive probe Oxyphor2P (Oxy2P) (Fig. 3i)^24^. Intravascular oxygen saturation (SO₂) was estimated from PO₂ using the Hill equation with coefficients specific to mice, as previously described^36^.

Analysis of arterial oxygenation revealed no significant differences between the sexes in either the pial or the diving arteries (Fig. 3j–3k). These results indicate that oxygen delivery to the cortical microvasculature is comparable between sexes at 15 months of age.

In contrast, the venous network showed clear sex-dependent differences. Venous PO₂ was significantly higher in females compared to males in both pial veins (45.16 ± 2.25 vs. 33.78 ± 1.58 mmHg; p = 0.0007) and diving veins (43.30 ± 1.09 vs. 34.08 ± 1.09 mmHg; p = 0.0081– Fig. 3l). Consistent with these findings, venous oxygen saturation was also significantly elevated in females relative to males in both pial (0.56 ± 0.03 vs. 0.38 ± 0.03; p = 0.0010) and diving veins (0.53 ± 0.01 vs. 0.39 ± 0.02; p = 0.0057–Fig. 3m).

Using arterial and venous oxygen estimates, we calculated OEF by pairing the median arterial and venous PO₂ values to estimate the oxygen concentration (cO₂). Female mice showed significantly lower OEF than males, consistent with the elevated venous oxygenation observed in the female microvasculature. This difference was significant in both pial vessels (p = 0.0009) and diving vessels (p = 0.0030; Fig. 3n).

To examine whether cortical perfusion was associated with oxygen extraction, we modeled OEF as a function of CBF and sex (Fig. 3o). The overall model was significant (F(3,7) = 6.46, p = 0.020, R² = 0.73). While CBF alone did not significantly predict OEF, a trend toward a sex-dependent interaction was observed (CBF*sex: β = 0.00246 ± 0.00126, p = 0.091). Specifically, higher CBF was associated with reduced OEF in females but with increased OEF in males.

Collectively, these findings suggest that the elevated cortical CBF observed in female mice at 15 months is accompanied by reduced oxygen extraction, consistent with a sex-dependent uncoupling between cerebral perfusion and oxygen utilization. Together with the stronger CTH-MTT coupling observed in females, these results point to altered capillary flow distribution, rather than macrovascular tone or vessel structure, as a key contributor to sex differences in cerebral oxygen delivery during aging. We next asked whether these microvascular differences were accompanied by molecular signatures of altered brain homeostasis.

### Reduced white matter volume is observed at 15 months in female mice

To assess whether structural differences accompanied the observed vascular and metabolic alterations, we performed ex vivo MRI volumetric analysis on a subset of brains (N = 4 per sex) collected at 15 months of age. Region segmentation allowed quantification of white matter (WM) and several subcortical structures, including the hippocampus (HC), caudoputamen (CP), thalamus (TH), and somatosensory cortex (SMS) (Fig. 1d). To avoid any confounding factor associated with the cranial window placed on the left hemisphere, we only analyzed the SMS of the right hemisphere. Hemispheric volumes revealed no significant differences between left and right hemispheres across regions (Fig. 1e-g).

Between sexes, total WM volume was modestly larger in males than in females for the whole brain (Est. = −0.818 ± 0.345 mm³, *p* = 0.0374, d = –2.577; Fig. 1e), whereas hemisphere-specific comparisons did not reach statistical significance (left: *p* = 0.142; right: *p* = 0.281). In contrast, we detected no sex differences in the volumes of the somatosensory cortex, hippocampus, caudoputamen, or thalamus.

### Spatially resolved transcriptomics reveals region-specific sex differences in the aging brain

After finding sex-dependent differences in cerebral hemodynamics, oxygen extraction, and WM volume, we asked whether these functional changes were accompanied by underlying transcriptional differences in brain tissue. In a separate age-matched cohort, we processed the left hemisphere for spatial transcriptomics using 10x Visium (N = 2 female, N = 2 male), generating transcriptome-wide expression profiles with spatial resolution of 55 µm per spot spot across regions segmented from H&E staining and manually identified using the Allen Mouse Atlas.

To identify broad transcriptional signatures that vary across tissue spatial contexts, we applied MEFISTO to the spatial RNA-seq data. MEFISTO models gene expression jointly with spatial coordinates and reduces the dataset to a smaller number of latent factors^37,38^. In this setting, that is useful because sex differences are unlikely to be driven by single genes alone, but rather by coordinated transcriptional programs that vary across anatomical regions. MEFISTO therefore provides a way to capture overall spatially structured expression changes first, and then test whether those latent programs differ between females and males.

We identified four interpretable latent factors with clear spatial and anatomical structure (Fig. 4). The representative maps showed that each factor occupied distinct tissue domains rather than diffuse gradients (Fig. 4a-d). The corresponding weight plots indicated that the factors were driven by different positively and negatively regulated gene sets, supporting the interpretation that they reflect distinct transcriptional programs rather than redundant axes of variation (Fig. 4e-h).

**Figure 4.**
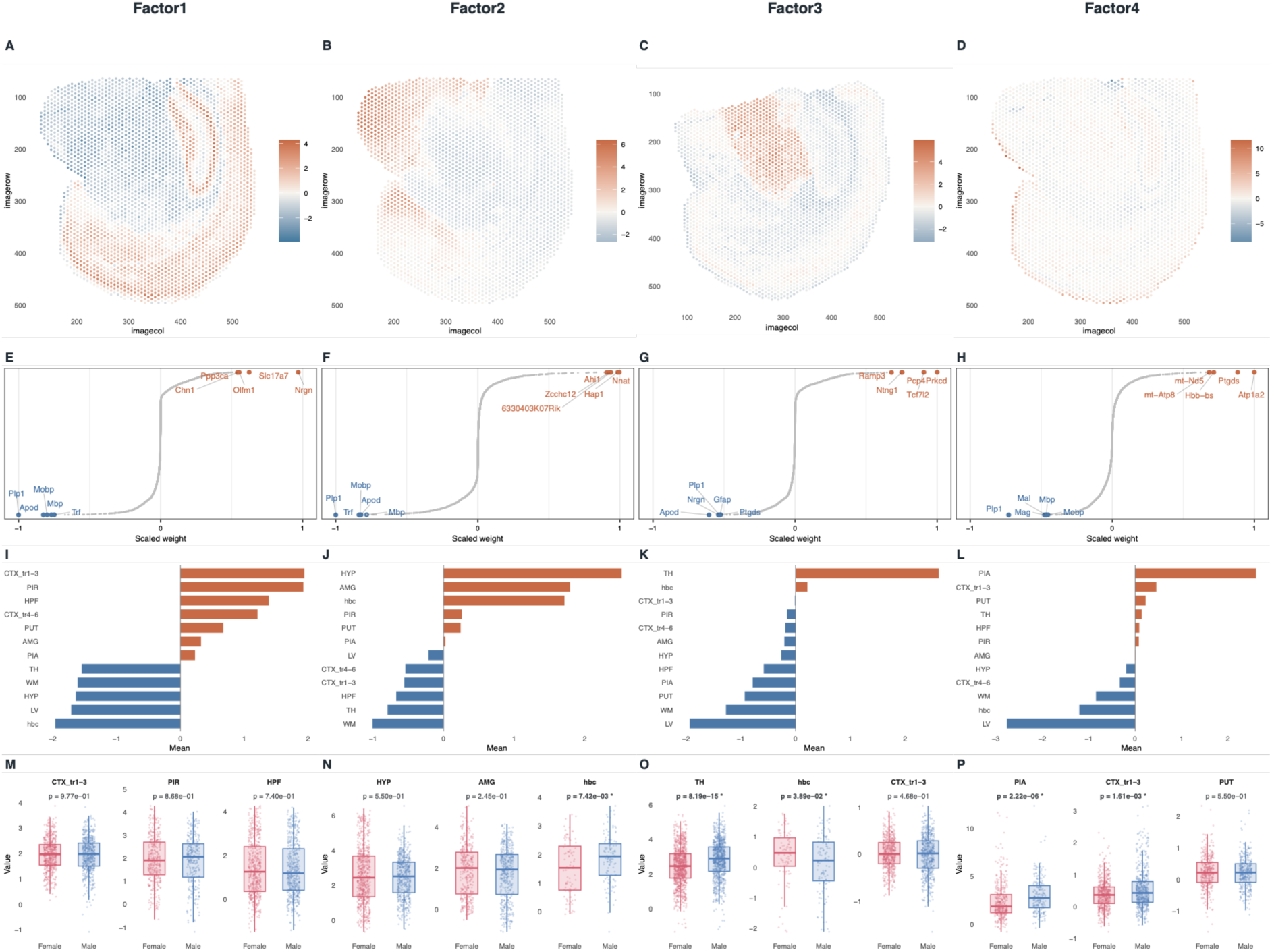
Spatial MEFISTO factors highlight anatomically structured latent programs and selected sex-associated differences. (A-D) Representative spatial maps for Factors 1-4, shown for the single tissue section that most clearly displayed the spatial pattern of each factor. Warmer colors indicate higher factor values and cooler colors indicate lower factor values within each panel. (E-H) Top positive and negative gene weights for each factor, illustrating the genes contributing most strongly to the positive and negative ends of the latent axis. (I-L) Mean factor values across annotated anatomical regions, summarizing the regional structure captured by each factor. (M-P) Spot-level factor distributions for the three regions with the highest positive mean values for each factor, shown separately for females and males. P values indicate spot-level female-versus-male Wilcoxon tests within each region, and significant comparisons (p < 0.05) are marked with an asterisk. Together, these panels show that Factors 1-4 capture distinct spatial programs with clear anatomical structure, while also indicating regional sex differences.

The factors primarily captured anatomical organization of the tissue (Fig. 4i-l). Factor 1 represented a strong cortex– and hippocampus-associated program opposed to white matter-like structure. Factor 2 was enriched in hypothalamic (HYP), amygdala-associated (AMG), and medial habénula (MHb) regions, consistent with a more restricted regional neuronal program. Factor 3 showed the clearest thalamus-associated (TH) pattern. Factor 4 was most strongly associated with the most superficial cortical region, assigned as PIA, and nearby boundary-like regions. Thus, the dominant signal recovered a region-specific signal. This regional factorization provided a useful framework for examining sex differences within biologically relevant spatial contexts (Fig. 4m-p). Together, these panels show that Factors 1-4 capture distinct spatial programs with clear anatomical structure, while also indicating regional sex differences.

Rather than comparing females and males across all spots pooled together, we compared spot-level factor values within the regions in which each factor was most strongly expressed. This showed that some factor-region combinations had shifted female and male distributions, whereas others remained largely overlapping. Factor 1 showed no significant sex differences in its top-associated regions. Factors 2 and 3 both showed significant sex differences in MHb, and Factor 3 additionally showed a difference in TH. Factor 4 showed differences in the upper cortical (CTX_tr1-3) and PIA. In contrast, not all top-ranked regions for each factor showed comparable separation between sexes. Therefore, sex differences were confined to specific latent programs and regions.

To characterize these regional sex differences at the gene level, we performed differential expression analysis within each region. DEG number varied substantially across regions, from a single gene in the lateral ventricle to over 400 in WM and TH, the two regions with both the largest number and the most statistically significant sex-dependent genes (Fig. 5a and 5b). Across regions, sex-chromosome genes (*Xist,* higher in females; *Ddx3y, Eif2s3y,* higher in males) were consistently the top differentially expressed genes, confirming correct sex assignment (Fig. 5b). Comparing DEGs across regions revealed that most genes were region-specific, with a smaller set shared across multiple regions (Fig. 5c-h).

**Figure 5.**
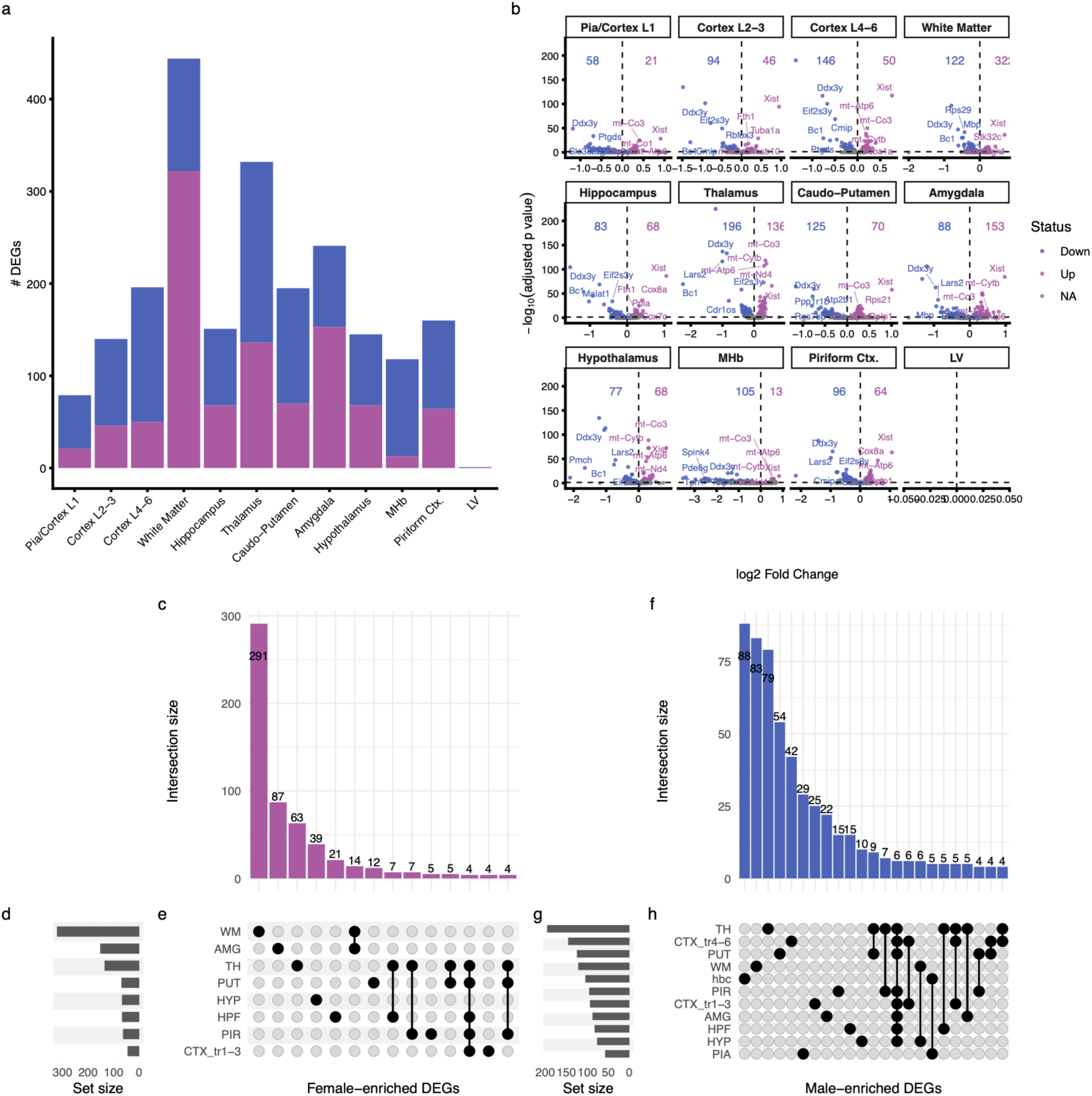
Region-specific and shared sex-dependent transcriptional differences across brain regions. (**a**) Total number of differentially expressed genes (DEGs; FDR < 0.05) per brain region, separated into female-enriched (magenta) and male-enriched (blue) genes. **(b)** Volcano plots for each brain region showing differential gene expression between sexes. Each point represents a gene, plotted by log2 fold change (female vs. male) and −log10(p-adjusted (Bonferroni)); selected top DEGs are labeled, and numbers indicate the total count of female-enriched (magenta) and male-enriched (blue) genes per region. **(c–e)** UpSet plot summarizing the overlap of female-enriched DEGs across brain regions. Intersection size (c) indicates the number of genes shared by each specific combination of regions. Set size (d) indicates the total number of female-enriched DEGs per region. The dot matrix (e) indicates which regions comprise each intersection. **(f-h)** As in (c-e), for male-enriched DEGs.

Among this shared set, gene ontology (GO) enrichment identified two dominant, recurring programs per sex and per region. Cytoplasmic translation and ribosome biogenesis were enriched among female-higher genes, strongest in the thalamus and caudoputamen, while synapse organization and neurotransmitter transport dominated male-enriched genes, most prominent in white matter, thalamus, and hippocampus (Fig. 6a and 6b). White matter showed the largest and most significant sex-dependent transcriptional differences of any region, but with a distinct directional profile relative to other regions. Cytoplasmic translation and ribosome biogenesis were enriched among male-higher genes (p.adj = 1.5×10^−^), the opposite pattern seen elsewhere, while synaptic vesicle cycle and membrane potential regulation genes were enriched among female-higher genes, also opposite relative to other regions. Male-higher genes were additionally enriched for axonogenesis (p.adj = 0.027), driven in part by the core myelin gene *Mbp*, though myelination and axon ensheathment did not reach significance as standalone GO categories, offering a candidate molecular correlate of the reduced white matter volume observed in females by ex vivo MRI.

**Figure 6.**
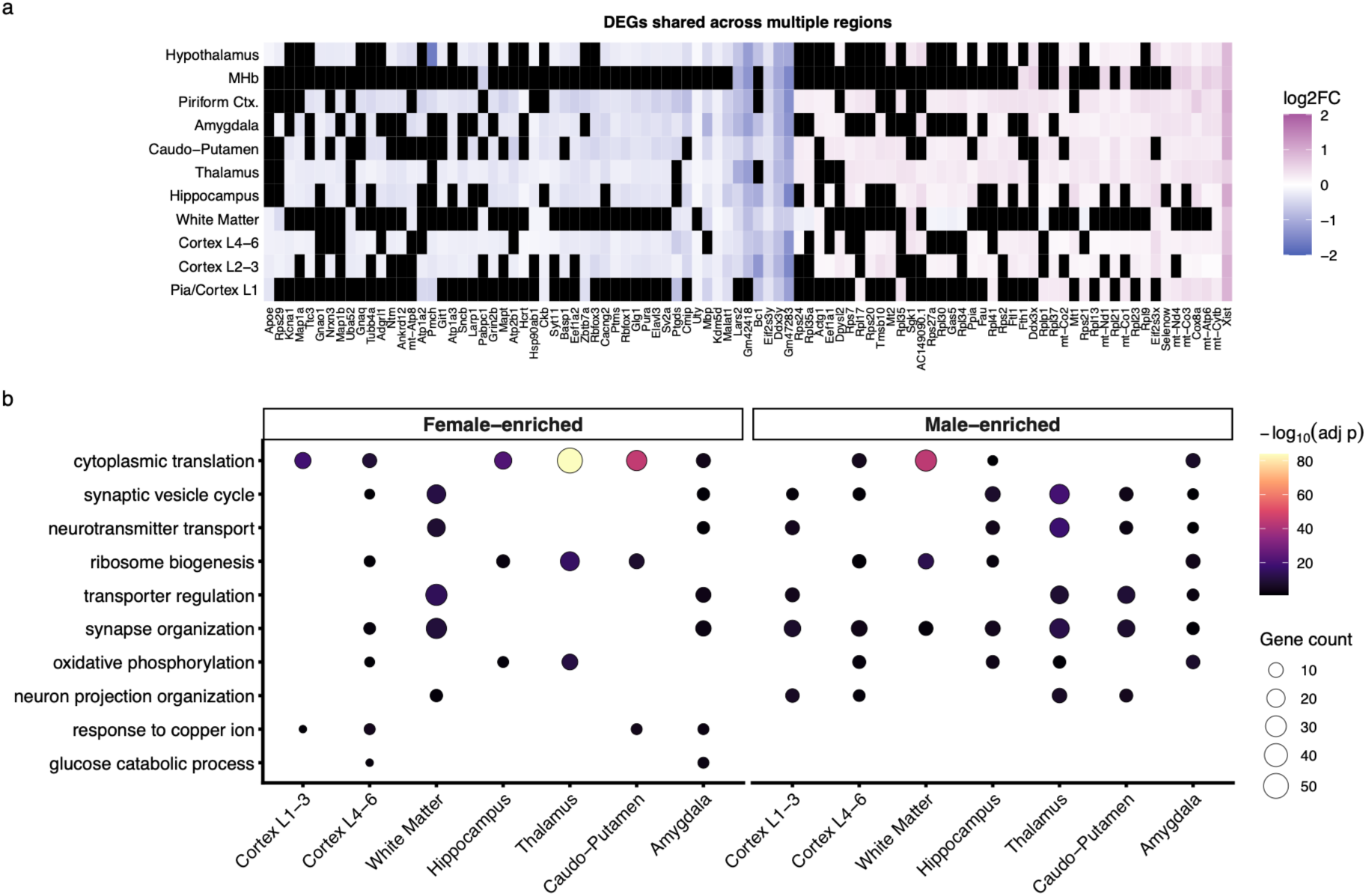
Shared and pathway-level sex-dependent transcriptional signatures. **a**. Heatmap of DEGs shared across five or more brain regions. Color scale represents log2 fold change (female vs. male); magenta indicates higher expression in females, blue indicates higher expression in males, and black indicates a non-significant difference for that gene in a given region. Genes (columns) are ordered by direction (female-enriched, left; male-enriched, right) and by the number of regions in which they are differentially expressed; regions (rows) are ordered by hierarchical clustering. **b.** GO enrichment analysis of differentially expressed genes, showing selected terms enriched among female– and male-enriched genes across brain regions. Dot color indicates statistical significance (−log10 adjusted p-value) and dot size indicates the number of genes contributing to each term.

We also observed a distinct, region-specific signal in the PIA region, where males showed gene enrichment for leukocyte migration and differentiation. TH and PIA were also among the regions where MEFISTO identified significant latent-factor sex differences, though the two analyses were not tested for shared underlying genes and should not be treated as an independent line of evidence pointing to the same regions rather than confirmed convergent biology.

Because our in vivo functional measurements were confined to the cortex and its pial vasculature, we examined the transcriptional signature in more detail. In both cortical layers segmented (L2-3 and L4-6), the same pattern seen elsewhere in the brain held. Synapse organization, neurotransmitter transport, and related neuronal terms were the dominant male signature (p.adj = 8.3×10⁻ in L2-3), while cytoplasmic translation and ribosome biogenesis dominated the female signature (p.adj = 1.1×10⁻¹ in L2-3, 4.3×10⁻¹ in L4-6). PIA region showed a qualitatively different pattern, consistent with its distinct vascular and meningeal composition. Male genes were enriched for smooth muscle cell proliferation, leukocyte migration, and response to steroid hormones (p.adj < 0.001), while female-higher genes showed the same translation/ribosome signature throughout the brain, though driven by a smaller gene set than in the cortical layers. This anatomically coherent distinction suggests that the most superficial cortical transcriptional sex difference plausibly reflects vascular/immune or perivascular cell populations rather than neuronal tissue, consistent with it being the tissue that later contains the pial vessels examined by optical imaging techniques.

Overall, we show that spatially coherent transcriptional programs define major anatomical structures, and that sex differences are not distributed uniformly across the brain but are instead concentrated in specific regions and organized around distinct translational, synaptic, and immune-related gene programs. Because our in vivo functional measurements were confined to the cortex and its pial vasculature.

### Vessel-associated astrocytic reactivity and increased capillary diameter variability distinguish the cortex of 15-month-old female mice

We next examined whether our findings were accompanied by detectable differences in vascular and glial markers at the tissue level. We performed immunohistochemistry on brains not used for ex vivo MRI, using the hemisphere that had undergone in vivo imaging.

Capillary diameter was quantified from CD31-labeled microvessels using VasoMetrics, measuring multiple capillary segments per mouse within predefined regions (cortex, hippocampus; Fig. 7a). For each mouse and region, we calculated the mean capillary diameter and its standard deviation (SD), the latter reflecting within-capillary diameter variability, using an averaged value per mouse. Mean capillary diameter did not differ significantly between sexes in cortex or hippocampus (Fig. 7b). Capillary diameter variability (SD) also did not differ significantly between sexes in either region, though hippocampus showed a large, non-significant effect (diff. = 0.150, F = 0.945, M = 0.795, p = 0.102, d = 1.274), while cortex showed a negligible difference.

**Figure 7.**
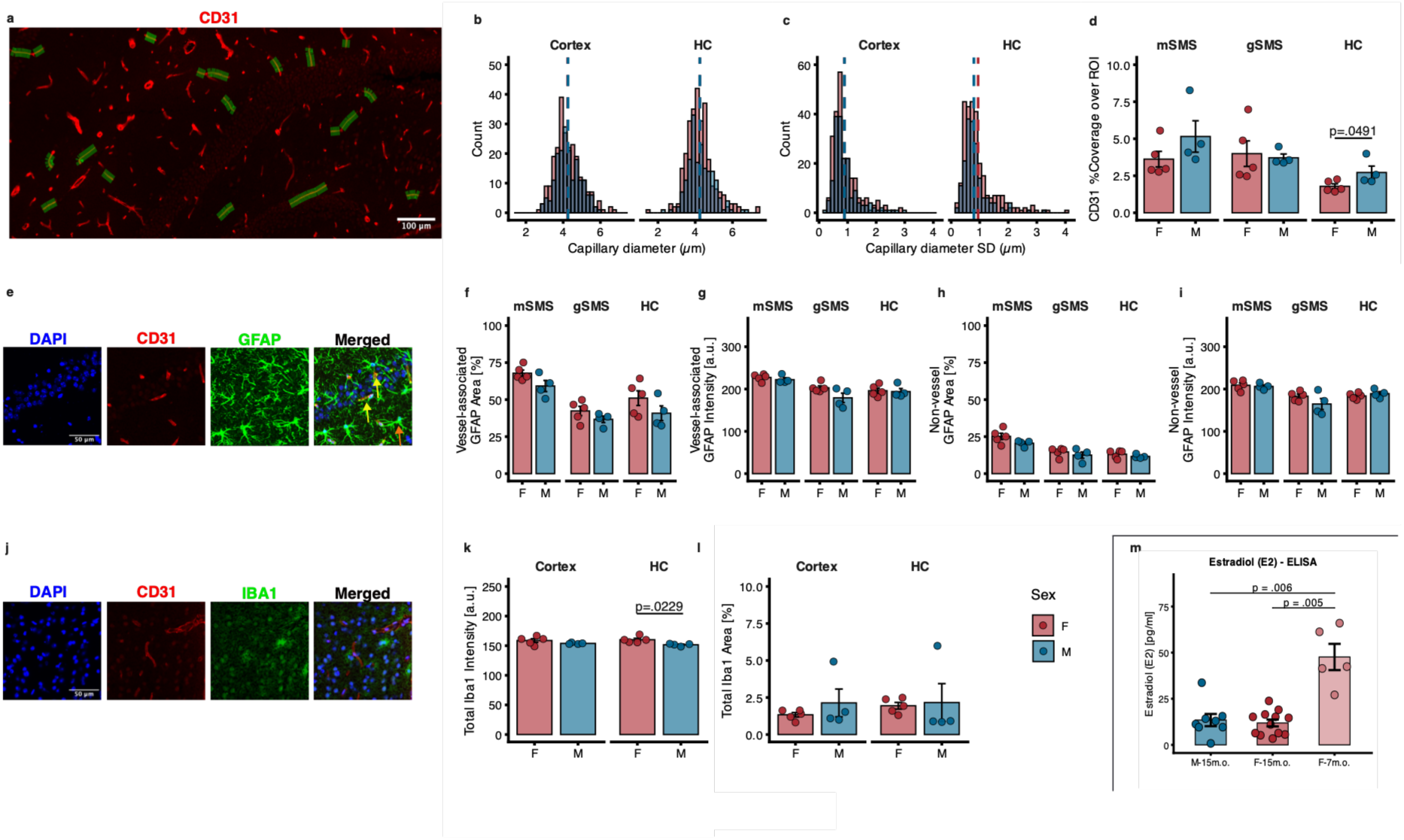
Sex differences in vascular, astrocytic, and microglial markers across brain regions. **a.** Representative image of CD31-positive vasculature (red) with example diameter measurements using vasometrics (green overlays).**b.** Distribution of capillary diameter in cortex and hippocampus (HC), showing comparable distributions between sexes. **c.** Distribution of capillary diameter variability (standard deviation), indicating similar heterogeneity across groups. **d.** Quantification of CD31 coverage (% area over ROI) across mSMS, gSMS, and HC, showing a significant sex difference in HC (p = 0.0491). **e.** Representative immunofluorescence images of DAPI (blue), CD31 (red), GFAP (green), and merged signal. Arrows indicate vessel-associated astrocytic processes. Scale bar: 30 µm **f–g.** Quantification of vessel-associated GFAP: **(f)** area fraction (%) and **(g)** mean intensity (a.u.). A non-significant trend toward higher vessel-associated GFAP in females was observed, most pronounced for area fraction in mSMS (d = 1.4110, p = 0.0762) and for intensity in gSMS (d = 1.429, p = 0.0641). **(h–i)** Quantification of non-vessel-associated GFAP: (h) area fraction (%) and **(i)** mean intensity (a.u.). No significant differences were observed between sexes in either measure, with the largest non-significant effects in mSMS area fraction (d = 1.213, p = 0.108) and gSMS intensity (d = 0.979, p = 0.213). **j**. Representative images of DAPI (blue), CD31 (red), Iba1 (green), and merged signal. Scale bar: 30 µm. **k.** Total Iba1 intensity (a.u.), showing increased signal in the hippocampus of females. **l.** Total Iba1 area (%), showing no significant differences between sexes. **m.** Plasma estradiol (E2) levels, measured by ELISA, were significantly higher in 7-month-old females than in both 15-month-old females (p = .005) and 15-month-old males (p = .006), with no difference between 15-month-old males and females (Kruskal-Wallis test followed by Dunn test for multiple comparisons). Error bars: mean ± SEM. F: female, M: male, ROI: region of interest, mSMS: molecular layer of the somatosensory area, gSMS: granular layer of the somatosensory area, HC: hippocampus, GFAP: glial fibrillary acidic protein, Iba1: ionized calcium-binding molecule 1.

For further immunohistochemistry analysis, we subdivided the somatosensory cortex (SMS) into a molecular layer (mSMS, corresponding to cortical layer 1) and a granular layer (gSMS, corresponding to layers 2/3-4), based on DAPI nuclear density/cytoarchitecture. Regions of interest (ROIs) were manually drawn in ImageJ/FIJI by an observer blinded to sex. Quantification of CD31 vascular coverage revealed a significant sex difference in the hippocampus, with greater coverage in males than in females (p = 0.0491, d = –1.470), with no difference observed in mSMS and gSMS (Fig. 7d).

To examine glial responses associated with the vasculature, we quantified GFAP immunoreactivity separately in vessel-associated and non-vessel-associated compartments (Fig. 7e). Vessel associated GFAP area fraction showed a non-significant trend toward higher values in females, most pronounced in mSMS (p = 0.0762, d = 1.411), with smaller, non-significant differences in HC and gSMS (Fig. 7f). Vessel associated GFAP mean intensity showed a similar non-significant trend in gSMS (p = 0.0641, d = 1.429), with no difference in HC or mSMS (Fig. 7g).

By contrast, non-vessel-associated GFAP showed no significant sex differences in either area fraction or intensity in any examined region. This pattern, with larger effect sizes concentrated in the vessel-associated compartment and minimal differences outside it, suggests that astrocytic changes in females may be preferentially localized to peri-vascular processes rather than reflecting a broader, tissue-wise astrocytic difference.

We next quantified microglial Iba1 immunoreactivity (Fig. 7j). Total Iba1 intensity was significantly higher in females in the hippocampus (p = 0.0229, d = 1.992), with no significant difference in the cortex (Fig. 7k). Total Iba1-positive area did not differ between groups.

Together, these histological findings indicate that the hemodynamic changes observed in female mice are accompanied by discrete structural and glial correlates at the tissue level.

### Estradiol decline confirms reproductive aging in the 15-month cohort

Because reproductive senescence in female mice is associated with declining circulating estrogen^39^, and because human menopause has been linked to changes in cerebral perfusion and metabolic vulnerability^5^, we measured plasma estradiol (E2) to confirm the hormonal aging status of our 15-month cohort relative to plasma samples from an independent cohort of younger reference group with the same background. E2 was significantly higher in 7-months-old females than in either 15-month-old females (p = 0.0047) or 15-month-old males (p = 0.0061), with no difference between 15-month-old males and females (Fig. 7m). Although female rodents typically show higher corticosterone than males^40^, this difference depends substantially on estradiol^41^, and its absence here is consistent with the low circulating estradiol in our 15-month females (p = 0.606; Fig.S4a). This confirms that female mice in our 15-month cohort had undergone the expected age-related decline in circulating estradiol, consistent with reproductive senescence, and by 15 months, circulating estradiol no longer distinguished the sexes.

Given that the regional differences could also arise from systemic factors acting on the brain through the circulation, we next examined whether vascular matrix remodeling could account for the convergence of PI and RI, the reduced hippocampal vascular coverage, and the trend toward higher vessel-associated GFAP in females. Circulating matrix metalloproteinase-9 (MMP-9), which degrades the vascular basement membrane and tight-junction proteins, has been linked to blood-brain barrier (BBB) disruption and to arterial stiffening^42^. We didn’t see any difference (p = 0.898; FigS4b). The absence of a sex difference in circulating corticosterone is consistent with human studies, in which baseline serum cortisol does not differ by sex even though its associations with brain structure and metabolism are sex-dependent^19^, suggesting that a null difference in circulating glucocorticoid levels does not rule out sex-specific downstream effects on brain physiology.

### Sex-dependent differences in spatial navigation emerge during middle age

The same cohort of mice underwent longitudinal behavioral testing at the 13-, 14-, and 15-month time points, spanning the in vivo imaging sessions described above. Consistent with previous reports of sex-specific behavioral alterations during middle age, including increased anxiety-like behavior and reduced locomotor activity in females^43^, we assessed spatial learning and behavioral adaptation using the Barnes maze with a similar paradigm each month (Fig. 8a).

**Figure 8.**
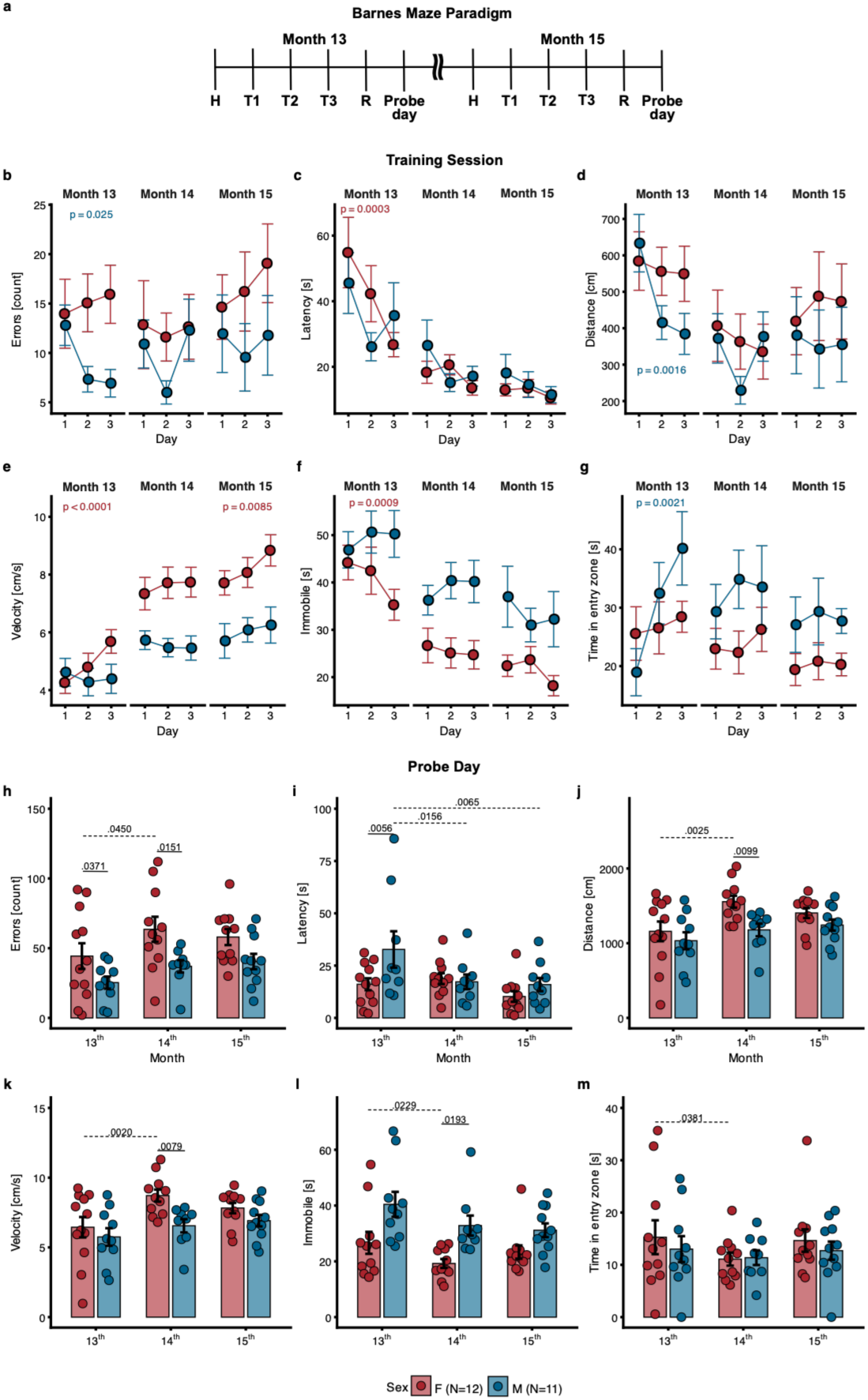
Sex– and Age-Dependent Behavioral Profiles in Barnes Maze Performance. **a**. Paradigm for the Barnes maze applied to the study. **(b–g)** Behavioral performance across training sessions in Months 13-15. Three sessions per day were averaged. Measures include **(b)** errors, (**c)** latency, (**d)** traveled distance, **(e)** velocity, **(f)** time spent immobile, and **(g)** time spent in the entry zone. Linear mixed-effects models (LMMs) were used to assess session-dependent changes (slopes) within each sex and differences between sexes. **(h–m)** Probe trial outcomes across age, including **(h)** errors, (**i)** latency, **(j)** traveled distance, **(k)** velocity, **(l)** time spent immobile, and **(m)** time spent in the entry zone. Individual data points represent single animals. Linear mixed-effects models (LMMs) were used to assess the effects of sex, age, and their interaction. F, female; M, male; bpm, beats per minute. Error bars: mean ± s.e.m. Statistical results are provided in the supplementary material.

During training, several sex-specific patterns emerged. In Month 13, male mice showed a significant reduction in errors across sessions (Fig. 8b), whereas females did not. We detected no significant sex differences in slopes and observed no session-dependent effects in Months 14 or 15. In contrast, latency to reach the escape zone decreased significantly across sessions in females during Month 13 (Fig. 8c), but not in males. Distance traveled during training also decreased significantly across sessions in males during Month 13 (Fig. 8d), but not in females, suggesting improved task efficiency early in training.

Females also showed changes in activity-related measures. The within-session slope of movement velocity, reflecting the rate of change across trials, differed significantly between sexes during Months 13 and 15, with females showing a steeper increase in velocity across sessions than males (Fig. 8e). Immobility time decreased during Month 13 (Fig. 8f), along with changes in time spent in the entry zone (Fig. 8g). Significant sex differences in slopes for velocity and immobility during Month 13 further support distinct early training adaptations between sexes.

On probe day (Fig. 8h-m), females made more spatial errors than males, and sex differences were significant at 13 and 14 months (Fig. 8h; 13 months, d = 1.250, p = 0.0371; 14 months, d = 1.613, p = 0.0105). These large effect sizes indicate a robust sex-dependent difference in spatial accuracy during middle age.

Because females also showed altered activity-related behaviors, with significantly greater distance traveled, higher velocity, and reduced immobility than males specifically at 14 months (Fig. 8j-l; d = 1.338, p = 0.0099; d = 1.374, p = 0.0079; d = –1.082, p = 0.0193, respectively), we next asked whether these differences could account for the elevated error rates, ruling out poorer task performance as the explanation. Notably, latency to the target zone did not increase in females and was even reduced relative to males at 13 months (Fig. 8i; d = –1.373, p = 0.0056), indicating that females reached the escape area more efficiently. Time spent in the entry zone did not differ significantly between sexes at any month (Fig. 8m), though females showed a significant within-sex decrease from 13 to 14 months (d = 0.886, p = 0.0381). Despite reaching the target more quickly and efficiently, females committed more spatial errors, suggesting altered search strategies rather than delayed task completion.

To determine whether these sex differences reflected impaired acquisition or retention, we next examined performance during training. Both sexes improved progressively across training sessions, and their escape probability curves largely overlapped, supporting comparable task acquisition (Supplementary Fig. S3a).

We then asked whether increased spatial errors were associated with locomotor-related behavioral states; we analyzed the relationship between errors and immobility time across ages and sexes (Supplementary Fig. S3b). At 13 months, errors were negatively correlated with immobility in both females (r = −0.74, BH-adjusted p = 0.011) and males (r = −0.90, BH-adjusted p = 0.002), indicating that animals with less immobility tended to show higher error rates. This correlation persisted in males at 14 and 15 months (r = −0.90, p = 0.003; r = −0.75, p = 0.012) but not in females (p = 0.119, p = 0.134), suggesting a more persistent coupling between locomotor activity and error rate in males across middle age.

Finally, to assess whether spatial errors reflected memory accuracy during probe testing, we correlated errors with time spent in the goal quadrant (Supplementary Fig. S3c). This relationship was significant only in females at 14 months (r = ∼0.79, BH-adjusted p = 0.024); no other age-by-sex comparison reached significance. This limited age– and sex-specific association suggests that the error metric’s relationship to spatial memory accuracy is not uniform and should therefore be interpreted cautiously rather than as a general index of memory performance.

Together, these findings indicate that middle-aged female mice show a sex-specific reduction in spatial accuracy that is not due to impaired learning, reduced locomotor capacity, or increased immobility. Instead, it reflects a sex-specific shift in exploratory strategy during spatial navigation.

## Discussion

This study examines whether sex differences in cerebral hemodynamics emerge in middle age and, if so, at what level of the vascular hierarchy they arise. First, cortical CBF was elevated in female mice at 15 months, despite convergence of earlier sex differences in PI and RI and the absence of detectable sex differences in pial vessel diameter or its short-term variability. Second, CTH scaled more steeply with MTT in females than in males, consistent with a more passive, less actively regulated capillary network. This was accompanied by reduced oxygen extraction driven specifically by the venous, capillary-level compartment rather than by any deficit in arterial oxygen delivery. Third, spatial transcriptomics revealed that sex differences were not uniform across the brain or uniformly female-associated, but organized into distinct, region-specific programs. Females showed a consistent translational and metabolic signature across regions, alongside synaptic, myelin, and leukocyte-migration programs concentrated in males in specific regions, including WM and the most superficial cortical region (PIA). These transcriptional differences were accompanied by measurable reductions in WM volume in females, elevated microglial Iba1 and reduced vascular coverage in the female hippocampus, and a shift in spatial navigation strategy in female mice that occurred without impaired learning or delayed goal reaching. Notably, these differences were not accompanied by sex differences in plasma markers selected to test the most plausible systemic explanations in this cohort (stress and vascular matrix remodeling), which favors a regional rather than a systemic origin. Estradiol was lower in the 15-month female cohort than in a younger reference group, consistent with the expected age-related decline in reproductive hormone status.

Sex differences in cerebral perfusion have shown inconsistent patterns across the human aging and menopause literature, with some studies reporting reduced CBF during middle age, particularly during the menopause transition, similar to male levels^3^, other finding no significant sex– or menopause-related differences in cerebrovascular reactivity^4^, and others describing elevated regional perfusion postmenopause as a compensatory response to declining brain volume and metabolism^5^. Against this mixed background, we found that cortical CBF was elevated in female mice at 15 months, at an age when earlier sex differences in PI and RI had already converged, indicating that this later divergence in bulk flow does not simply track macrovascular pulsatility and instead points toward disturbances in the microcirculation as its likely source.

Theoretical models of capillary flow predict that oxygen availability and extraction efficiency depend not only on CBF, but on the degree of CTH, with greater heterogeneity limiting extraction efficiency at a given flow rate^8^. Consistent with this framework, we found that CTH scaled more steeply with MTT in females than in males, indicative of a more passive, less actively regulated capillary network, and that this was accompanied by reduced oxygen extraction driven specifically by the venous, capillary-level compartment rather than by any deficit in arterial oxygen delivery. We previously observed similar scaling in an animal model of cerebral amyloidosis^13^ and a model of water intoxication^44^. In the latter, we associated astrocytic endfoot swelling with disrupted microvascular flow regulation. This raises the possibility that estrogen-dependent astrocyte reactivity, shown to emerge at midlife in female mice (12-14 months) following loss of ovarian hormone signaling^45^, could similarly compromise capillary flow regulation, though this link remains speculative and was not directly tested here. Future studies using models of induced ovarian hormone loss, such as ovariectomy, combined with in vivo capillary flow imaging, would be needed to directly test this hypothesis.

A recent study performing spatial transcriptomics of the aging brain has identified WM as a region of disproportionate age-related transcriptional change, associated with a stronger neuroinflammatory response in females specifically in this region across aging in mice^16^. In our dataset, obtained at a single midlife timepoint, WM also showed the largest and most significant sex-dependent transcriptional differences in any region, though the specific signature differed. Rather than an inflammatory axis, the dominant distinction was between a male-associated myelin and axonogenesis program and a female-associated translational and ribosomal program. Notably, none of the five plasma markers of stress, blood-brain barrier integrity, or humoral immunity differed between sexes, indicating that these differences were regionally confined rather than reflecting a systemic inflammatory or endocrine state. This transcriptional signature was accompanied by reduced WM volume in females on ex vivo MRI, offering a candidate structural correlate, altough individual-gene rather than pathway-level significance of myelin association limits our interpretation. Because mice do not undergo menopause, but rather a gradual reproductive senescence accompanied by reduced systemic estrogen, our 15-month female cohort cannot be mapped precisely onto human menopausal staging. Human diffusion imaging studies have found that WM organization in women differs by menopausal stage in a non-monotonic way. Pre– and postmenopausal women show denser, more organized WM than age-matched men, while perimenopausal women show a transient shift toward a more male-like profile^17^. Whether our findings correspond to this perimenopausal-like transitional pattern, or instead reflect a distinct, mouse-specific trajectory of reproductive aging, remains an open question that our cross-sectional hormone data alone cannot resolve.

Hippocampal transcriptomic studies in mice have reported sexually divergent, complement-enriched neuroinflammatory gene induction with aging^14^, with sex differences emerging at old age; females show enriched senescent and disease-associated microglial signatures linked to a shift toward glycolytic metabolism in aged female hippocampal microglia^15^. Although our microglial analysis did not include morphology to better characterize inflammatory activation, the elevated hippocampal Iba1 intensity observed in females is consistent with this evidence that microglial aging is sexually dimorphic in this region. This microglial metabolic shift is distinct from the bulk-tissue translational signature we observed transcriptionally, but both point to metabolic reprogramming. Further studies assessing hippocampal hemodynamics are needed to understand how this sexual dimorphism in microglial activation relates to disturbances in hippocampal flow dynamics and tissue oxygenation.

Female mice showed more spatial errors during the Barnes maze probe testing at 13 and 14 months, with a similar, non-significant trend at 15 months, despite reaching the escape zone as efficiently as, or more efficiently than, males, a pattern better explained by an altered exploratory or search strategy than by a general, consistently replicated impairment in spatial memory. Sex differences in rodent spatial navigation and memory tasks during midlife vary considerably across studies and testing paradigms, rather than following a consistent pattern. In the Morris water maze, females have been reported to swim faster than males without finding the platform earlier, suggesting a less directed navigational strategy rather than a memory deficit^46^. In another Barnes maze study spanning 4 to 14 months of age, sex did not significantly interact with age for either training performance or probe-trial errors, though a modest overall main effect of sex was observed^47^. In a separate study extending to 24 months, aged females showed impaired spatial and reversal learning relative to males, while male cognitive flexibility remained preserved^48^. Together, these findings suggest that the female-specific increase in spatial errors we observed reflects a change in navigational strategy rather than a general spatial memory deficit, consistent with a broader literature in which sex differences in rodent spatial cognition are inconsistent across paradigms and emerge when present, though distinct behavioral mechanisms rather than a uniform pattern of impairment.

## Limitations

Several limitations should be considered when interpreting our findings. Sample sizes were modest and unevenly distributed across modalities. Spatial transcriptomic comparisons were based on two animals per sex; while differential expression was tested at spot-level resolution, the biological replication for these findings rests on this smaller animal-level sample, and results should be interpreted as candidate signatures for validation in larger cohorts. Hemodynamic measurements were restricted to the cortex and pial vasculature, and 2P imaging was performed at 15 months; therefore, we cannot determine when capillary-level differences emerge or whether regions showing the largest transcriptional differences, such as WM and the thalamus, share them. Lastly, our hormonal data are cross-sectional, and without estrous cycle staging we cannot establish whether 15-month females were in a stable reproductive-senescent state or an ongoing transition, which limits comparison with staged human menopausal cohorts. Finally, a single plasma corticosterone measurement reflects hypothalamic–pituitary–adrenal axis activity at the time of sampling rather than cumulative stress exposure, BBB integrity was not assessed directly by parenchymal IgG or fibrinogen extravasation.

## Conclusion

In this cohort of middle-aged mice, sex differences in cerebral hemodynamics were absent or transient at the macrovascular pial level but emerged at the capillary network level. Female mice showed elevated cortical CBF, steeper coupling between CTH and MTT, and reduced OEF, without changes in input oxygen delivery, suggesting that changes in oxygen availability are limited to the capillary or venous compartments. These changes coincided with region-specific transcriptional programs, reduced WM volume, regional glial and vascular differences in the hippocampus, and an altered spatial search strategy. They were observed at an age when circulating estradiol in females had declined to levels similar to those in males, and without sex differences in circulating markers of stress, blood-brain barrier integrity, or vascular matrix remodeling. Because some studies have reported elevated CBF in postmenopausal women, and we observed it here in aged females with low circulating estradiol, yet it coincided with reduced oxygen extraction, bulk CBF may not reliably reflect tissue oxygen delivery in this setting. Future studies of reproductive aging should therefore assess capillary regulation directly, including the roles of ovarian hormone signaling and astrocityic function.

## Author Contributions

E.G.J. conceptualized the study, performed validation, formal analysis, and investigation, provided resources, curated the data, prepared the visualizations, wrote the original draft, reviewed and edited the manuscript, supervised and administered the project, and acquired funding. B.W. contributed to the methodology, validation, and investigation, including surgical preparation, laser speckle data acquisition, and training of the mice, and reviewed and edited the manuscript. T.L. contributed to conceptualization, methodology, and validation, including MRI acquisition and training of the mice, and reviewed and edited the manuscript. S.M. contributed to the methodology and validation, including surgeries, training of the mice, and two-photon imaging, and reviewed and edited the manuscript. P.M.R. contributed to the methodology and developed software for the oxygen measurements, and reviewed and edited the manuscript. D.P. contributed to the methodology, developed the laser speckle analysis software, reviewed the results, and reviewed and edited the manuscript. B.H. contributed to conceptualization and methodology, including study design and the analysis of pCASL data, and reviewed and edited the manuscript. O.B. performed formal analysis of the imaging data, curated the data, prepared visualizations, and reviewed and edited the manuscript. M.E.K. and S.V. contributed to the methodology through the synthesis and calibration of Oxy2P, and reviewed and edited the manuscript. F.M. performed formal analysis of behavioral data, curated the data, and reviewed and edited the manuscript. A.K. contributed to the methodology through the establishment of the Bessel beam and reviewed and edited the manuscript. L.L. performed the formal analysis of the spatial transcriptomics data, wrote the corresponding Methods section of the original draft, and reviewed and edited the manuscript. J.J. performed the MEFISTO analysis of the spatial transcriptomics data, contributed to the corresponding Results section in the original draft, and reviewed and edited the manuscript. I.P.P. performed ELISA experiments and reviewed the manuscript. R.W.K. and C.S.S. performed MRI scans and analysis, and reviewed and edited the manuscript. M.F.J. performed analysis of LSCI. V.D. curated the data, prepared visualizations, contributed to the original draft, and reviewed and edited the manuscript. L.Ø. contributed to conceptualization, provided resources, and reviewed and edited the manuscript.

## Supporting information

Statistical analysis tables

## Acknowledgments

We thank John Giblin for support with setting up the software for the behavioral stalls, Susanne Smith Christensen for laboratory technical support, Anna Bay for immunohistochemistry and ELISA support, and Camille Szeto for support with the immunohistochemistry analysis.

