## Supplementary material for "Sex shapes capillary flows and oxygen extraction in the middle-aged mouse brain"

Supplementary materials

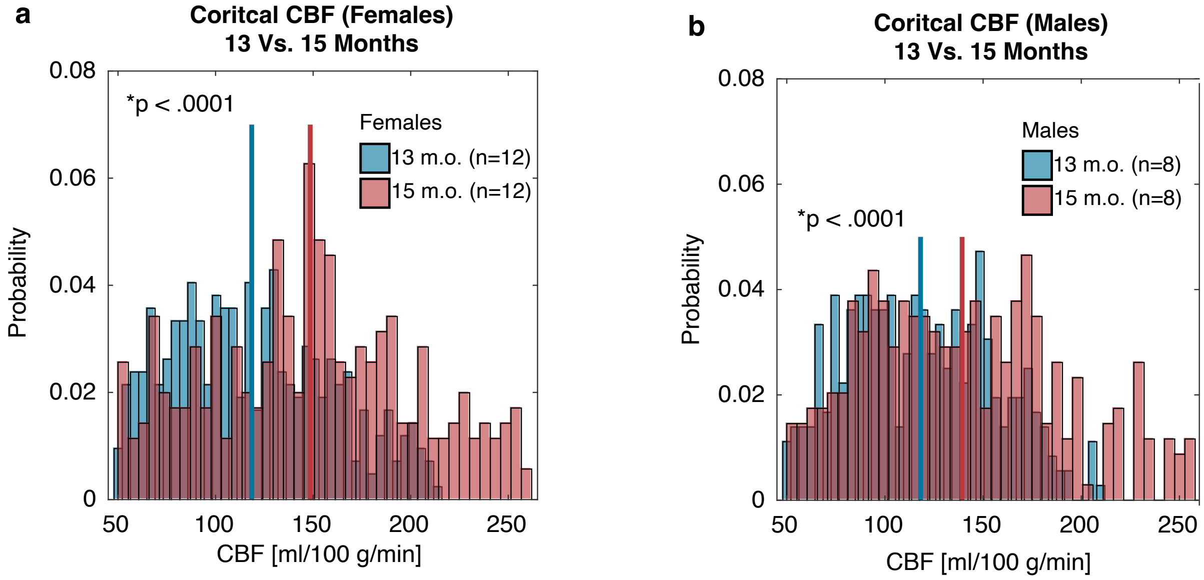


**Figure S1. Increased in CBF is observed in female and male mice from 13 to 15 months of age. a.** Histogram of pixel-wise CBF values in the cortex of female mice at 13 months (blue; n = 12) and 15 months (red; n = 12). Mean CBF increased significantly from 118.53 ± 40.89 (blue line) to 148.49 ± 52.68 ml/100 g/min (red line, $p < 0.0001$). **b.** Histogram of pixel-wise CBF values in the cortex of male mice at 13 months (blue; n = 8) and 15 months (red; n = 8). Mean CBF increased significantly from 118.14 ± 37.05 (blue line) to 139.31 ± 49.65 ml/100 g/min (red line, $p < 0.0001$). Statistical analysis: Permutation tests (100,000 permutations) with t-tests of means were used to compare 13 vs. 15 months within each sex. Error bars: mean ± SEM.

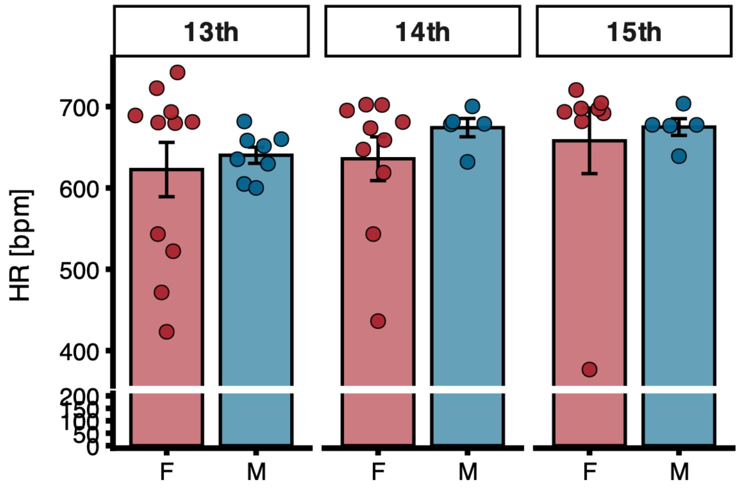


**Supplementary Figure S2.** Heart rate derived from LSCI.

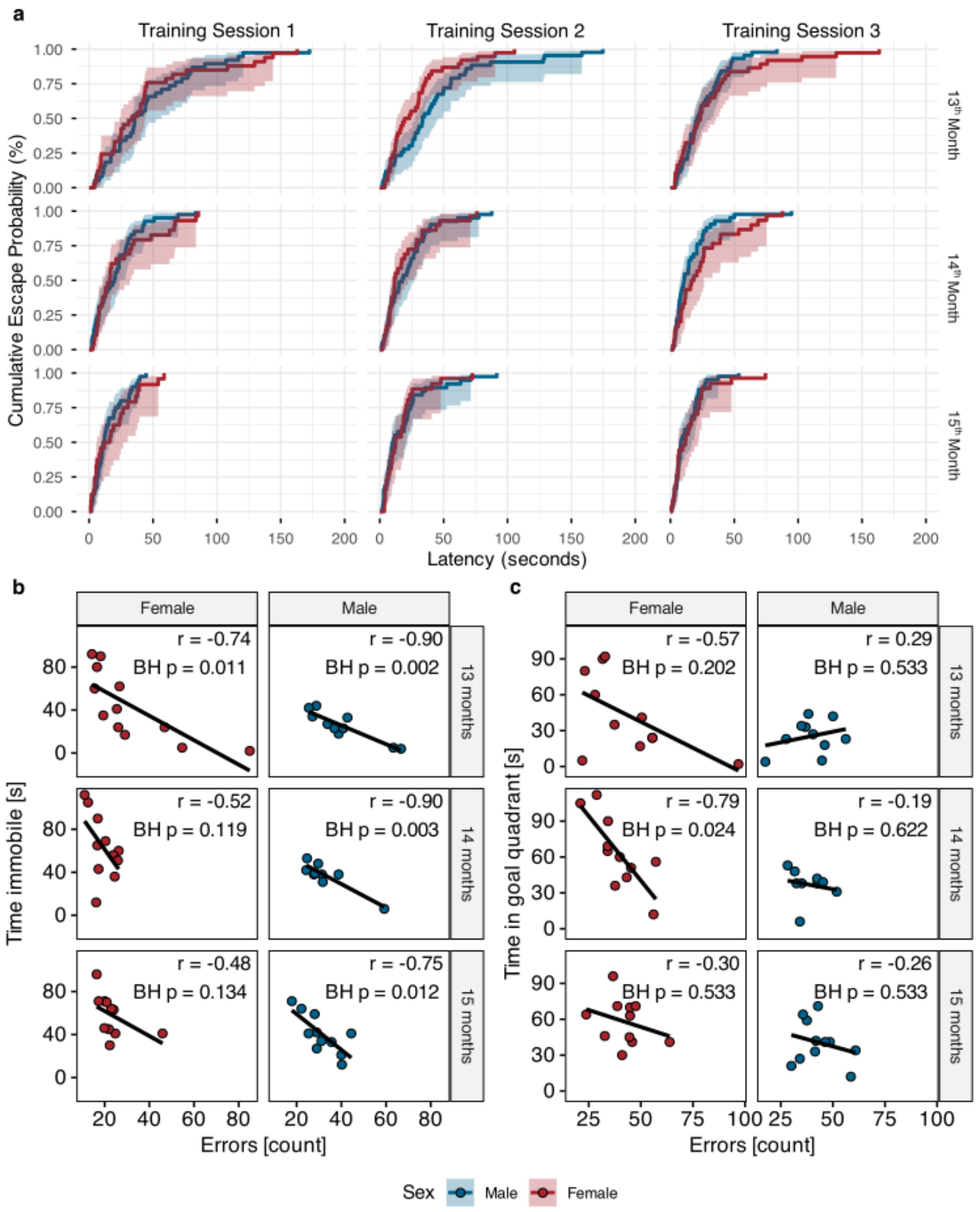


**Supplementary Figure S3. (a)** Cumulative plot depicting escape probability during Barnes maze training sessions 1–3 at 13, 14, and 15 months of age. Curves represent Kaplan–Meier estimates of escape latency distributions for females (red) and males (blue). Shaded areas indicate confidence intervals. **(b)** Correlation between errors and time immobile during probe testing, stratified by sex and age. Pearson correlation coefficients (r) and Benjamini–Hochberg (BH)-adjusted p values are shown within each panel.**(c)** Correlation between errors and time spent in the goal quadrant during probe testing. A significant negative correlation was observed only in females at 14 months (BH-adjusted $p < 0.05$), indicating that higher error levels were associated with reduced spatial accuracy in this group. No significant associations were detected in males. Data points represent individual animals. Lines denote linear regression fits.

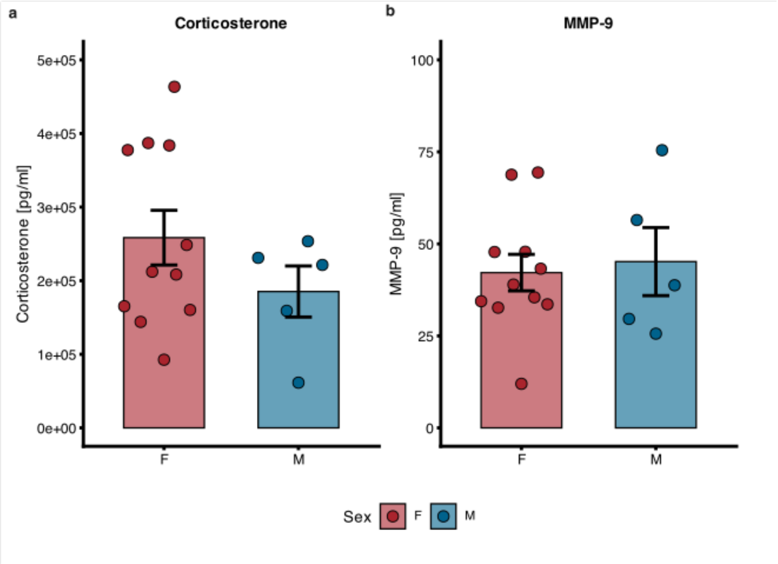


**Supplementary Figure S4. Serum corticosterone and MMP-9 levels do not differ by sex.** Bar plots show mean ± SEM concentrations of corticosterone (a) and MMP-9 (b) in plasma from female (F) and male (M) mice. Individual points represent individual animals. No significant sex differences were detected for any marker (Mann-Whitney U test).
